# Natural predator-associated chemosignals evoke distinct spatiotemporal patterns of risk assessment and vomeronasal population responses

**DOI:** 10.64898/2026.10.30.763133

**Authors:** Jinxin Wang, Varun Haran, Julian P. Meeks

## Abstract

Natural predator-associated chemosignals elicit diverse defensive behaviors, yet it remains unclear how complex natural cues shape the spatiotemporal organization of risk assessment and activate vomeronasal sensory neurons (VSNs) and downstream circuits in the mammalian accessory olfactory system. Here, we investigated odor-dependent behavioral organization and VSN sensory representations using three-dimensional behavioral tracking, integrative machine-learning analysis, volumetric calcium imaging, and whole-brain FosTRAP2 mapping. We developed a pose-based integrative machine-learning analysis pipeline (PIMAP) to identify behavioral signatures based on posture, spatial exploration, state transitions, and temporal dynamics. We found that common behavioral states were expressed in multiple odor conditions, but their expression rates, spatial positions, and temporal sequencing distinguished responses to snake, tortoise, lizard, and mouse fecal cues. Snake fecal cues elicited sustained direct investigation and distinct patterns of spatial exploration, behavioral transitions, and grooming in both two-chamber and multiple-choice assays. Volumetric calcium imaging in the vomeronasal epithelium revealed distinct patterns of VSN activation by different reptilian cues. Whole-brain microscopy of FosTRAP2 transgenic mice indicated potential differential activity patterns in AOS-associated circuits following snake and tortoise exposure. Furthermore, Trpc2 deletion altered snake-associated behavioral signatures, while odor-discriminative behavioral patterns remained detectable. Together, these findings show that reptilian predator cues activate distinct AOS components and bias exploration patterns towards threat assessment and vigilance.

## Introduction

Predator-associated chemosignals provide critical information about potential danger, allowing animals to detect and evaluate potential threats before direct encounters (Apfelbach et al., 2005; Blanchard et al., 2011). In mammals, the accessory olfactory system (AOS) detects biologically relevant chemical signals, including conspecific pheromones and interspecific kairomones (Munger et al., 2009; Holy, 2018; Mohrhardt et al., 2018). Previous studies implicate the AOS in detecting predator-derived chemosignals and shaping defensive responses, while also demonstrating contributions from parallel olfactory pathways (Masini et al., 2010; Papes et al., 2010; Pérez-Gómez et al., 2015). These findings establish an important role for the AOS in predator chemosensation. However, how sensory detection through this system relates to the behavioral responses elicited by complex natural predator-associated cues remains unclear.

Natural predator feces contain complex chemical mixtures that may recruit diverse populations of vomeronasal sensory neurons (VSNs) (Haran et al., 2026), raising questions about how the vomeronasal organ (VNO) represents chemically heterogeneous predator-associated chemical cues. VSNs express diverse receptor families, including V1Rs and V2Rs, that detect distinct classes of chemical signals (Dulac and Torello, 2003; Isogai et al., 2011; Mohrhardt et al., 2018). Previous studies have identified predator-derived proteins and small-molecule ligands, including fecal bile acids, that activate distinct VSN populations (Papes et al., 2010; Doyle et al., 2016; Wong et al., 2020). Population-level studies further demonstrated distinct, partially overlapping VSN responses to sulfated steroids and urine samples from mice of different sexes and strains, supporting population-level representations of defined ligands and complex chemosignals (Turaga and Holy, 2012; Fu et al., 2015). Natural fecal chemosignals may convey multiple biological features of the source animal, including species, diet, and gut microbial composition (Haran et al., 2026). Although individual vomeronasal ligand classes have been characterized, it remains unclear whether snake fecal cues evoke VSN population activity patterns distinguishable from those elicited by other reptile fecal cues or conspecific mouse feces.

Chemosensory information detected by VSNs is transmitted to the accessory olfactory bulb (AOB), where it is processed before reaching the medial amygdala (MeA) and other limbic regions (Meeks et al., 2010; Keshavarzi et al., 2015; Holy, 2018). The MeA and downstream hypothalamic circuits contribute to processing predator-associated cues and organizing defensive responses, including approach-avoidance behaviors (Gross and Canteras, 2012; Pérez-Gómez et al., 2015; Miller et al., 2019). Previous studies have demonstrated that predator-related information from the main olfactory system (MOS) and AOS can converge in the ventromedial hypothalamus, where organized neuronal ensembles represent chemosensory information and predator-threat imminence (Pérez-Gómez et al., 2015; de Andrade Carvalho et al., 2020; Nguyen et al., 2024). Despite these advances, how complex natural predator chemosignals recruit distributed brain regions and how these responses relate to risk-assessment behavior remain unclear. Genetic disruption of *Trpc2* alters vomeronasal-dependent social and defensive behaviors, including responses to predator-derived cues (Leypold et al., 2002; Stowers et al., 2002; Papes et al., 2010). However, predator-associated signals can also engage parallel olfactory pathways (Masini et al., 2010). Thus, it remains unclear whether *Trpc2*-dependent signaling contributes to the postural and spatiotemporal organization of risk-assessment behavior elicited by natural predator-associated chemosignals.

Predator-associated chemosignals elicit diverse and multidimensional behavioral responses, ranging from cautious investigation, including flat-back approaches and stretch-sniffing, to avoidance, hiding, and altered non-defensive activities (Dielenberg et al., 2001; McGregor et al., 2004; Apfelbach et al., 2005; Papes et al., 2010; Blanchard et al., 2011; Nguyen et al., 2024). These responses also vary with stimulus intensity, exposure history, and environmental context (Staples et al., 2008; Takahashi, 2014). Conventional behavioral measurements, such as freezing, avoidance, and investigation duration, capture individual components but may overlook their coordinated spatial, postural, and temporal organization. Recent advances in markerless pose estimation, supervised behavioral classification, and unsupervised behavioral segmentation enable high-resolution quantification of movement, recurring postural patterns, and behavioral dynamics across different timescales (Berman, 2018; Mathis and Mathis, 2020; Saoud et al., 2024; Fazzari et al., 2025). Although these approaches provide increasingly detailed descriptions of naturalistic behavior, computationally identified states do not necessarily capture their biological significance, as similar movements may have different meanings depending on spatial location, environmental context, and preceding or subsequent behaviors. For example, direct odor investigation involves a characteristic head-down posture with the nose positioned near or in contact with the odor source, whereas similar postures elsewhere in the arena may reflect general exploration. Thus, integrating high-resolution behavioral measurements with postural and spatiotemporal information and ethologically grounded interpretation remains essential for characterizing the complex organization of predator-evoked risk-assessment behavior.

To address these questions, we integrated three-dimensional (3D) behavioral analysis, volumetric VSN calcium imaging, FosTRAP-based brain activity labeling, and genetic disruption of *Trpc2*. We first extended our previous hybrid machine-learning (ML) workflow (Wang et al., 2023) to develop a 3D pose-based integrative machine-learning analysis pipeline (PIMAP) for characterizing behavioral responses to natural reptilian fecal cues in a two-chamber risk-assessment task and a multiple-choice odor-investigation task. Next, we used objective-coupled planar illumination (OCPI) Ca²⁺ imaging to characterize VSN population responses to reptile and mouse fecal extracts (Wong et al., 2020). We also characterized behavioral responses in an independent FosTRAP2 cohort and used whole-brain serial two-photon tomography (STPT) to map regional neuronal labeling and explore its relationship to behavior. Finally, we compared wild-type (WT) and *Trpc2* knockout (KO) mice to examine the contribution of *Trpc2*-associated vomeronasal signaling to odor-elicited behavioral organization. Together, these experiments characterize peripheral sensory responses, brain-wide FosTRAP2 labeling, and multidimensional behavioral patterns associated with natural predator chemosignals, providing a foundation for investigating their central neural representations.

## Materials and Methods

### Animals

All animal procedures were approved by the University Committee on Animal Resources at the University of Rochester and followed the Guide for the Care and Use of Laboratory Animals. Mice were maintained on a 12-h light/dark cycle with ad libitum access to food and water. Behavioral experiments were performed during the dark cycle under dim red illumination. Wild-type (WT) C57BL/6J mice (The Jackson Laboratory, stock #000664) and *Trpc2* knockout (KO) *Trpc2^tm1Dlc^* mice (*Trpc2*^−/−^; stock #021208) were used for behavioral experiments (Stowers et al., 2002; Cansler et al., 2017). For VNO Ca²⁺ imaging, OMP-Cre mice (Omptm4(cre)Mom/J; stock #006668) (Li et al., 2004) were crossed with Ai96 mice (Gt(ROSA)26Sortm96(CAG-GCaMP6s)Hze/J; stock #028866) (Madisen et al., 2015) to express GCaMP6s in VSNs, as described previously (Wong et al., 2020). For brain-wide activity labeling, mice carrying the FosTRAP2 allele (Fos-2A-iCreERT2) and homozygous for the Cre-dependent Ai9 reporter allele (Rosa26-lox-STOP-lox-tdTomato) were used (Madisen et al., 2010; DeNardo et al., 2019; Ramirez et al., 2019). Experiments were performed on adult mice aged 8-16 weeks. Animal numbers and sex distributions are provided in the corresponding experiments and figure legends.

### Fecal sample preparation and odor stimuli

Reptile fecal samples were collected from Seneca Park Zoo (Rochester, NY) and grouped according to the diet of the source animals. The mouse-fed carnivore group included Eastern massasauga rattlesnake (*Sistrurus catenatus*) (Snake1) and Dumeril’s ground boa (Snake2) (*Acrantophis dumerili*); the insect-fed omnivore group included Henkel’s gecko (Lizard1) (*Uroplatus henkeli*) and great plated lizard (Lizard2) (*Broadleysaurus major*); and the salad-fed herbivore group included pancake tortoise (Tortoise1) (*Malacochersus tornieri*) and green iguana (*Iguana iguana*) (Iguana1). Female mouse feces were collected from C57BL/6 mice and used as a conspecific odor stimulus. Aqueous fecal extracts were prepared as described previously (Wang et al., 2023; Haran et al., 2026). Female mouse fecal extract was prepared using the same procedure. Distilled water (dH_2_O) was used as the control stimulus in the behavioral experiments. 2,4,5-Trimethylthiazole (TMT) (Sigma-Aldrich, catalog #219185) was used as an aversive odor reference. TMT was diluted 1:9 in mineral oil to produce a 10% solution, as described previously (Saito et al., 2017; Wang et al., 2023).

### Behavioral experiments

Behavioral experiments were performed during the dark cycle under dim red illumination. Mouse behavior was recorded using four synchronized FLIR Blackfly cameras (BFS-U3-16S2C-CS, USB 3.1) positioned to obtain right, left, bottom, and top views of the arena. Videos were acquired at 27 frames/s. Odor presentation and arena handling followed procedures established in our previous risk-assessment study (Wang et al., 2023). For fecal odor presentation, 3D-printed artificial fecal particles (∼1 cm long, elliptical shape) were immersed in the corresponding fecal extract overnight. Particles (n=5-7) were placed in a Petri dish and kept moist during testing. Separate Petri dishes were used for each odor to minimize cross-contamination.

The two-chamber risk-assessment task was modified from our previously established risk-assessment paradigm (Wang et al., 2023). The arena consisted of a hiding chamber [2 in (L) × 8 in (W) × 8 in (H)] connected to an exploration chamber [15 in (L) × 8 in (W) × 8 in (H)] by an open doorway in a transparent divider. The odor-containing Petri dish was fixed at the distal end of the exploration chamber, and mice could move freely between chambers. Snake1, Lizard1, Tortoise1, female mouse feces, dH₂O, and 10% TMT were used as the Snake, Lizard, Tortoise, Mouse, Control, and TMT conditions, respectively. Each odor exposure lasted 5 min in Cohorts 1 and 3. For Cohort 2, mice were exposed to odors for 10 min, and the first 5 min of behavior were used for analysis. Between two experimental mice, fecal odor-containing particles (n=5-7) were returned to their corresponding extract tubes to maintain moisture and refresh the stimuli before being placed back into their respective Petri dishes for the next mouse. Between animals, the behavioral arena was cleaned with 75% ethanol to minimize residual olfactory cues from previously tested mice. Three independent experimental cohorts of mice were used. Cohort 1 included 12 WT mice (6 males and 6 females) tested with all six odor conditions. Mice were habituated to the arena with dH₂O on Days 1-3, followed by a dH₂O Control session on Day 4. Mice were then exposed to Lizard (Day 5), female mouse feces (Day 6), Snake (Day 7), Tortoise (Day 8), and TMT (Day 9). Cohort 2 included independent groups exposed to Control (n=11; 6 males and 5 females), Snake (n=15; 7 males and 8 females), and Tortoise (n=12; 6 males and 6 females) and was used for behavioral analysis associated with FosTRAP2 odor exposure and labeling, followed by STPT acquisition for subsequent brain-wide activity analysis. Mice were habituated on Days 1-2 and exposed to the assigned odor on Day 3 for FosTRAP2 labeling. Cohort 3 included 9 WT and 9 *Trpc2* KO mice (3 males and 6 females per genotype) and was tested with the same six odor conditions as Cohort 1.

To examine behavioral responses when different natural odors were encountered simultaneously, 24 mice were tested in the multiple-choice odor-investigation task using a square arena [18 in (L) × 18 in (W) × 6 in (H)]. Mice freely explored the arena for 20 min while behavior was recorded using the same synchronized multi-camera system. Mice were habituated to the arena on Days 1 and 2, during which Petri dishes containing dH₂O were placed in all four corners. On Day 3, Snake1, Lizard1, Tortoise1, and female mouse feces were assigned randomly to the four corners of the testing arena for the initial session, and these assignments were maintained for subsequent mice. Between two experimental mice, fecal odor-containing particles (n=5-7) were returned to their corresponding extract tubes to maintain moisture and refresh the stimuli before being placed back into their respective Petri dishes for the next mouse. Maintaining these odor locations across animals minimized repeated handling and the potential for cross-contamination during stimulus refreshing. The two-day habituation and single odor-exposure design were intended to minimize novelty-related exploration and avoid spatial preference due to repeated odor-location associations. Between animals, the behavioral arena was cleaned with 75% ethanol to minimize residual olfactory cues from previously tested mice.

### FreiPose 3D pose estimation

3D mouse pose was reconstructed from synchronized multi-camera recordings using FreiPose (Schneider et al., 2022). Camera intrinsic and extrinsic parameters were estimated and calibrated using FreiCalib, and calibration quality was assessed by FreiCalib-derived reprojection error. For the two-chamber experiments, Cohort 1, Cohort 2, and Cohort 3 models had reprojection errors of 4.76, 5.51, and 4.56 pixels, respectively. The multiple-choice model had a calibration reprojection error of 4.10 pixels.

Twelve anatomical landmarks were tracked, including the nose, left and right eyes, left and right ears, left and right front paws, left and right hind-paw ankles, left and right hind-paw tips, and the tail base. The models for Cohorts 1, 2, and 3 were trained using 426, 172, and 463 manually labeled frames, respectively. The multiple-choice model was trained using 528 manually labeled frames. Model performance was quantified using mean per-joint position error (MPJPE), defined as the mean 3D Euclidean distance between predicted and manually labeled landmark positions (Schneider et al., 2022). MPJPEs were 1.43, 2.08, and 2.64 mm for the two-chamber models of Cohorts 1, 2, and 3, respectively, and 3.51 mm for the multiple-choice model. Although the Petri dishes remained fixed during testing, their positions and geometry were determined independently for each video. Eight landmarks equally spaced around each dish circumference were manually labeled in three frames. Their reconstructed coordinates were averaged across frames and used to calculate the dish center and diameter, providing spatial references for dish-relative behavioral variables. To enable direct comparison across the three two-chamber cohorts, reconstructed spatial measurements were converted to physical units using the known 55-mm Petri-dish diameter as an internal reference. Cohort-specific scaling factors were applied to place reconstructed coordinates on a common physical scale. The multiple-choice dataset was analyzed independently and did not require additional scaling. Distances were expressed in millimeters, and velocities were expressed in mm/s after accounting for the 27-frames/s acquisition rate.

### Pose-based integrative machine-learning analysis pipeline

Behavior was quantified using the Pose-based Integrative Machine-learning Analysis Pipeline (PIMAP), an upgraded 3D version of our previously established layered hybrid ML-based workflow (Wang et al., 2023). The original workflow was based primarily on 2D positional measurements and integrated supervised Random Forest classification with Gaussian hidden Markov models (GHMM) to capture temporal behavioral structure. Here, we extended this framework to three dimensions by incorporating multi-camera FreiPose tracking (Schneider et al., 2022) and expanded the unsupervised behavioral analysis beyond the original Gaussian HMM approach using Keypoint-MoSeq (KPMS) and state-based hierarchical MoSeq (shMoSeq). KPMS identifies sub-second behavioral syllables directly from pose dynamics (Weinreb et al., 2024), whereas shMoSeq, using the output from KPMS, organizes sequences of these syllables into longer-timescale behavioral states (Weinreb et al., 2026). Together, these additions enabled PIMAP to capture behavioral organization across spatial, postural, and temporal scales beyond those represented in our original 2D workflow.

#### Supervised Random Forest behavioral classification

Following established supervised behavioral annotation approaches (Kabra et al., 2013) and our previous RF-based behavioral analysis workflow (Wang et al., 2023), separate Random Forest (RF) classifiers were trained for the two-chamber and multiple-choice behavioral tasks to identify interpretable behaviors from 3D pose-derived features. RF models were implemented using RandomForestClassifier in Scikit-learn (Pedregosa et al., 2011).

For the two-chamber task, ten videos from the three cohorts (seven 10-min and three 5-min recordings; 137,700 frames) were manually annotated frame-by-frame as ground truth. Five behavioral classes were defined: direct odor investigation, rearing, grooming, wall climbing, and exploration, which included all other behaviors. Direct odor investigation was defined as a head-down posture in which the mouse’s nose was positioned within the Petri dish and was either in direct contact with or in close proximity to the odor particles. Rearing was defined by elevation of the forequarters while supported by the hind limbs. Grooming included paw licking, face/head washing, and body grooming. Wall climbing required an elevated posture with at least one forepaw contacting the arena wall/divider. All remaining behaviors were annotated as exploration. The model used 121 FreiPose-derived postural features and 22 Petri-dish-relative positional features. For the multiple-choice task, three 20-min videos (97,200 frames) were manually annotated. The same behavioral definitions were used, except that direct odor investigation was separated into four classes according to the Petri dish being investigated, resulting in eight behavioral classes. The model used 117 FreiPose-derived postural and global positional features and 31 Petri-dish-relative positional features.

To incorporate local temporal information and improve RF classification, each behavioral feature was smoothed using rolling median filters with window sizes of 3, 5, 7, 11, 15, 30, and 60 frames, extending the temporal-feature strategy used in our previous workflow (Wang et al., 2023). The original and all smoothed features were combined for model training, yielding 1,144 predictors for the two-chamber model and 1,184 predictors for the multiple-choice model. Model performance was evaluated using 10-fold stratified cross-validation at the annotated-frame level (StratifiedKFold, shuffle=True, random_state=42) with out-of-fold predictions. The two-chamber model was evaluated on 137,700 manually annotated frames and achieved an accuracy of 0.977 and macro-averaged F1 of 0.966. The multiple-choice model was evaluated on 97,200 manually annotated frames and achieved an accuracy of 0.979 and macro-averaged F1 of 0.964.

#### Keypoint-MoSeq and shMoSeq

The 3D FreiPose coordinates were first fed into the Keypoint-MoSeq model (version 0.5.0), an unsupervised generative framework that links keypoint tracking to pose dynamics and segments continuous behavior into recurring sub-second behavioral syllables (Weinreb et al., 2024). Separate Keypoint-MoSeq models were fitted for the two-chamber and multiple-choice datasets. For both assays, pose dynamics were represented using 10 principal components, and the model was configured with the default setting of 100 syllables. The stickiness parameter kappa (k), which controls the tendency of the model to remain within the same syllable and thereby influences the temporal scale of behavioral segmentation (Weinreb et al., 2024), was adjusted separately for each dataset to achieve a median syllable duration of approximately 500 ms (∼13 frames). For the two-chamber dataset, k was set to 5 ×10⁷, and the model was fit for 550 iterations. For the multiple-choice dataset, k was set to 1,500, and the model was fit for 350 iterations.

To characterize behavioral organization at a longer timescale, the sequences of Keypoint-MoSeq syllables were further analyzed using state-based hierarchical MoSeq (shMoSeq; state-moseq, installed package version 0.0.0) (Weinreb et al., 2026). Separate shMoSeq models were fitted to the two-chamber and multiple-choice datasets. The number of higher-order states was set to 17 for both datasets, with trans_kappa=12000 and trans_beta=1. Emission parameters were set to emission_base_sigma=1 and emission_biases_sigma=1. Model optimization used 1,000 gradient-descent iterations with a learning rate of 5 × 10⁻³ and random_seed=0.

#### Integrative Gaussian HMM and final PIMAP state definition

To integrate supervised behavioral classification, unsupervised pose dynamics, and spatial information relative to the odor source, we extended our previously established layered HMM-based analytic strategy (Wang et al., 2023). Frame-aligned RF classifications, shMoSeq states, and selected Petri-dish-relative variables were used as inputs to a GHMM (hmmlearn.GaussianHMM). Input features were standardized before GHMM fitting. The spatial variables captured behaviorally relevant relationships between the animal and odor source, including nose height relative to the dish and head/nose distance from the dish in the two-chamber task, and nose-to-dish distances and directional information in the multiple-choice task. Candidate GHMMs with 10-30 hidden states were evaluated for model convergence, log-likelihood, and behavioral interpretability. Because the supervised RF classifications represented the most directly interpretable behavioral categories, confusion matrices comparing GHMM states with RF classes were used to determine whether candidate models preserved distinct behavioral categories while avoiding unnecessary state fragmentation. The hidden states generated by this integrative GHMM model were initially defined as PIMAP behavioral states.

Candidate PIMAP states were interpreted using synchronized video clips, 3D skeletal reconstructions, spatial distributions within the arena, and their relationships to the supervised and unsupervised model outputs. Behaviorally redundant states were consolidated only when their observed pose, movement pattern, and spatial context were sufficiently similar. The final two-chamber dataset, including three cohorts, comprised 24 PIMAP states used for downstream analyses of state occupancy, transitions, temporal dynamics, and odor-associated behavioral signatures. For the multiple-choice dataset, five PIMAP states were defined within each odor quadrant, allowing equivalent behavioral states to be compared across quadrants within the same animal. To assign behavioral states to individual quadrants, the animal’s body centroid was compared with the center of the arena, which was determined from manually labeled arena landmarks. The four quadrants were defined according to the body centroid’s position relative to the arena center. This assay complemented the single-odor two-chamber assay by characterizing behavioral responses when different natural chemosignals were simultaneously available.

### Behavioral feature extraction and linear discriminant analysis

Behavioral features were extracted from the final PIMAP state sequences to capture complementary aspects of behavioral organization, including state occupancy, bout structure, movement, state-transition probabilities. For temporal analysis, each recording was divided into two equal halves, and changes in behavioral features were calculated as the second-half value minus the first-half value. For the multiple-choice assay, “Entry with [behavior]” denotes the percentage of quadrant entries beginning with that behavior, whereas the corresponding count describes how many such entries occurred. State transition probabilities represent the likelihood of moving between two behavioral states. The two-chamber analysis generated 1,634 behavioral features from the 24 PIMAP states, whereas the multiple-choice analysis generated 144 features from the quadrant-specific PIMAP states. The complete feature sets and definitions are summarized in Supplementary Tables 1 and 2.

To compare behavioral responses across odor conditions in this high-dimensional feature space, we used linear discriminant analysis (LDA), following the multivariate strategy established in our previous behavioral-analysis workflow (Wang et al., 2023). LDA was implemented using *sklearn.discriminant_analysis.LinearDiscriminantAnalysis* from Scikit-learn (Pedregosa et al., 2011). A separate LDA model was fitted for each experimental cohort and for the multiple-choice dataset. For Cohort 3, WT and *Trpc2* KO mice were analyzed using separate LDA models. The resulting LDA projections were used to characterize behavioral differences within the fitted datasets and guide subsequent behavioral-signature discovery.

### Behavioral signature discovery and optimization

Behavioral signatures were identified using a multistep post-LDA workflow that extended the multivariate behavioral-analysis strategy established in our previous study (Wang et al., 2023):

1. Primary LD selection. For each target odor, Cohen’s *d* was calculated between the target and the pooled remaining odor groups along each LD. The LD with the largest absolute target-versus-rest Cohen’s *d* was defined as the primary LD.
2. LD-based feature ranking. Behavioral features were ranked according to the absolute magnitude of their weights on the primary LD, and the N_top features with the largest absolute weights, including both positively and negatively weighted features, were retained as candidate features.
3. Effect-size screening. Candidate features were screened using Cohen’s *d* for predefined target-versus-comparator contrasts. Features were retained only when the required comparisons exceeded the specified absolute Cohen’s *d* threshold. The comparator groups used to select each odor-associated signature set are specified in the corresponding figure legends.
4. Signature classification. Individual features passing the effect-size screen constituted the behavioral signatures for each target odor group. For each target odor’s signature set, a multiclass logistic regression model with L2 regularization (C=1.0) was trained on standardized features to classify the odor conditions. Model performance was evaluated using four-fold stratified group cross-validation repeated four times. This procedure was applied to all three two-chamber cohorts and the multiple-choice dataset. Out-of-fold predicted probabilities were averaged across repetitions to calculate target-class one-versus-rest receiver operating characteristic (ROC) curves and areas under the curve (AUCs).

To optimize signature size and discriminability, N_top and the minimum Cohen’s *d* threshold were systematically varied. For the two-chamber cohorts, N_top was tested at 25, 50, 75, 100, and 150, and the Cohen’s *d* threshold was varied from 0.3 to 0.8. Parameter combinations were evaluated by considering target-class cross-validated AUC together with the number of retained signatures. When classification performance was comparable, settings yielding fewer signatures were preferred, while larger signatures were retained only when further signature reduction resulted in a substantial decrease in AUC. A single parameter combination was applied to all target odors within each cohort to maintain a consistent screening criterion. Because the parameter optimization was performed separately for each cohort, the selected N_top and Cohen’s *d* thresholds were allowed to differ across cohorts. The selected settings were N_top=50 and |d| ≥ 0.4 for Cohort 1; N_top=25 and |d| ≥ 0.5 for Cohort 2; and N_top=50 and |d| ≥ 0.5 for Cohort 3. The same Cohort 3 settings were applied to WT and *Trpc2* KO mice to permit direct genotype comparison. For the multiple-choice dataset, N_top=10 and |d| ≥ 0.5 were used. The resulting cross-validated AUCs summarized the discriminability of the selected signatures within the analyzed datasets.

To examine the contribution of *Trpc2* signaling to WT-defined behavioral signatures, the signatures selected from Cohort 3 WT mice and the corresponding WT-trained logistic regression models were subsequently applied to *Trpc2* KO observations without additional signature selection or model retraining. This analysis evaluated whether the behavioral patterns used to discriminate odor conditions in WT mice were preserved in *Trpc2* KO mice. For signature-level genotype comparisons, WT and KO signatures were identified independently using the same three-criterion Cohen’s *d* screening procedure, requiring |d| ≥ 0.5 for the target odor versus Control, TMT, and pooled remaining natural odors. Signatures were classified as WT-only (selected exclusively in WT), shared (selected in both genotypes), or KO-only (selected exclusively in KO). For each feature selected as a signature in either genotype, Cohen’s *d* was calculated separately within both genotypes by comparing the target odor with the pooled remaining three natural odors, excluding Control and TMT. These effect sizes were used to characterize changes in magnitude and direction, such as preserved, enhanced, attenuated, lost, reversed, and emergent-in-KO.

### Volumetric VNO Ca²⁺ imaging

#### Fecal stimulus panel and stimulus presentation

The fecal sample sources used in the OCPI experiments are described in the “Fecal sample preparation and odor stimuli” section. Reptile fecal extracts from Snake1, Snake2, Lizard1, Lizard2, Tortoise1, and Iguana1 were tested at 1:300 and 1:1,000 dilutions, and female mouse fecal extract was tested at 1:1,000 as a conspecific comparison. Extracts were diluted in Ringer’s solution containing 115 mM NaCl, 5 mM KCl, 2 mM CaCl₂, 2 mM MgCl₂, 25 mM NaHCO₃, 10 mM HEPES, and 10 mM glucose. Ringer’s solution alone served as the negative control. Stimuli were applied for approximately 15 s using an air-pressure-operated reservoir and multibarrel perfusion system (AutoMate Scientific, Berkeley, CA, USA). Stimulus delivery was randomized and interleaved under computer control, following our established VNO imaging procedures (Turaga and Holy, 2012; Wong et al., 2020; Haran et al., 2026).

#### VNO preparation and OCPI imaging

Following deep isoflurane anesthesia and rapid decapitation, VNOs were dissected and the vomeronasal epithelium was carefully removed under a dissection microscope. The epithelium was mounted on nitrocellulose paper and placed in a custom imaging chamber. Volumetric GCaMP6s imaging was performed using a custom OCPI microscope with previously described refinements (Turaga and Holy, 2012; Wong et al., 2020; Haran et al., 2026). Custom software synchronized fluorescence imaging with randomized, interleaved stimulus delivery. Each image stack contained 51 optical sections spanning approximately 700 µm laterally, 250-400 µm axially, and ∼150 µm in depth, and stacks were acquired once every 3 s (∼0.33 Hz). Each stimulus was delivered during the acquisition of five consecutive stacks (∼15 s), with at least 10 stacks (≥30 s) between stimulus trials. At least three complete randomized, interleaved stimulus blocks were acquired for each analyzed preparation.

#### Analysis of volumetric VNO Ca²⁺ imaging

Volumetric imaging data were analyzed using custom MATLAB procedures established in our previous studies (Wong et al., 2020; Haran et al., 2026). Image stacks were aligned across time using repeated rigid registration supported by SimpleElastix (Marstal et al., 2016). Following registration, 3D regions of interest (ROIs) encompassing individual VSN somata were manually defined using a custom MATLAB interface. Mean voxel fluorescence was calculated for each ROI in every image stack, generating a fluorescence time series for each VSN. For each stimulus presentation, ΔF/F was calculated by subtracting the mean fluorescence of three consecutive prestimulus stacks from the mean fluorescence of three stacks acquired during stimulus delivery and dividing by the mean prestimulus fluorescence. Across-trial mean ΔF/F was then calculated for each ROI and stimulus. To control for fluorescence changes associated with valve switching, stimulus-evoked responses were compared with responses to Ringer’s control stimulation. ROIs were included in subsequent population analysis when at least one stimulus produced a positive mean ΔF/F response >10% and the across-trial stimulus response differed from Ringer’s control with *p* < 0.1 using a two-tailed Student’s *t* test. Normalized VSN response profiles were clustered using density-based merging (DBM) implemented in the MATLAB runUMAP package (Walther et al., 2009). Cluster parameters were selected by inspection of the resulting response structure and pairwise LDA, which was used to evaluate the separation between clustered VSN response profiles. Cluster separation was quantified using the discriminability index (Cohen’s *d*), with *d* > 1.64 used as an empirical criterion for satisfactory separation between response clusters. These clustered response profiles were used to characterize differences in VSN tuning across snake, lizard, tortoise/iguana, and mouse fecal cues.

### FosTRAP2 labeling and 3D behavioral recording during odor exposure

The Cohort 2 two-chamber risk-assessment task was performed to characterize behavioral responses during the odor exposures used for subsequent brain-wide activity labeling. FosTRAP2×Ai9 mice were assigned to Control, Snake1, or Tortoise1 groups. Following two days of habituation, mice were exposed to the assigned odor on Day 3 during the FosTRAP2 labeling session. FosTRAP2 provides permanent genetic access to neurons expressing Fos within a temporally restricted 4-hydroxytamoxifen (4-OHT)-dependent recombination window (Guenthner et al., 2013; DeNardo et al., 2019). 4-OHT was prepared fresh on the day of the experiment following our established TRAP procedure (Zuk et al., 2023). Briefly, 4-OHT was dissolved in ethanol, mixed with an equal volume of corn oil, and the ethanol was removed under vacuum to produce a final concentration of 10 mg/ml. Mice received 4-OHT by intraperitoneal injection at 50 mg/kg 20 min before odor exposure, allowing recovery from the injection procedure before behavioral testing. During the 5-min odor-exposure session, behavior was simultaneously recorded using the four-camera 3D tracking system and analyzed using the PIMAP workflow described above.

#### Tissue collection and serial two-photon tomography

Seven days after odor exposure and FosTRAP2 labeling, mice were deeply anesthetized and transcardially perfused with phosphate-buffered saline followed by 4% paraformaldehyde. Brains were removed and post-fixed in 4% paraformaldehyde before preparation for serial two-photon tomography (STPT). STPT acquisition and image analysis followed established whole-brain imaging workflow (Ragan et al., 2012; Poinsatte et al., 2019; Ramirez et al., 2025).

STPT was performed using a TissueCyte 1000 system (TissueVision, Inc., Newton, MA). Agarose-embedded brains were mounted on a magnetic stage within an imaging chamber containing phosphate buffer. STPT was performed as an automated block-face imaging procedure in which two-dimensional mosaic images were acquired in the coronal plane before removal of the imaged tissue with an integrated vibrating microtome (Ragan et al., 2012; Ramirez et al., 2025). Three optical planes were acquired at 25, 50, and 75 µm below the cut surface, followed by removal of a 75-µm physical section. The vibrating microtome was operated at 70 Hz with an advancement velocity of 0.5 mm/s. This procedure generated approximately 190 physical sections and 570 optical-section images spanning the rostrocaudal extent of each brain, with lateral pixel sampling of 0.875 µm/pixel and axial sampling intervals of 25 µm. Excitation and emission channels were selected to collect endogenous tdTomato fluorescence together with tissue autofluorescence required for anatomical registration.

#### Image processing and automated whole-brain FosTRAP2 quantification

Image processing followed the workflow described in previous whole-brain STPT studies (Poinsatte et al., 2019; Ramirez et al., 2023; Ramirez et al., 2025). Raw image tiles were trimmed, subjected to flat-field correction, and stitched into two-dimensional coronal mosaic images using Autostitcher software (TissueVision, Inc.). For each physical section, the three optical planes were combined to generate maximum-intensity projection images using custom MATLAB procedures. Image contrast was linearly adjusted to reduce intensity differences across images, following established processing workflow (Ramirez et al., 2025). Images used for subsequent classification and registration were downsampled to 1.5 µm/pixel. Whole-brain datasets were visually inspected for structural or imaging artifacts before automated analysis.

Automated quantification followed the established STPT informatics pipeline consisting of supervised ML-based pixel classification, registration of whole-brain image volumes to the Allen Mouse Brain Common Coordinate Framework version 3.0 (CCFv3), and automated quantification of fluorescent signal within anatomically defined brain regions (Poinsatte et al., 2019; Wang et al., 2020; Ramirez et al., 2023; Ramirez et al., 2025). FosTRAP2-associated tdTomato signal was identified using the Pixel Classification Workflow ilastik, which implements a pixel-wise Random Forest classifier (Berg et al., 2019). Representative images (n=20) from sample brains were included in the training dataset and manually labeled to distinguish tdTomato fluorescence within neuronal cell bodies (class 0, P0) and neuronal processes (class 3, P3). A single classifier was trained and applied consistently across all brains. Classification outputs were exported as separate scaled 8-bit probability maps for P0 and P3, in which a pixel intensity of 255 corresponded to a classification probability of 1.0 for the respective class. Probability maps were median filtered before subsequent analysis to reduce high-contrast noise artifacts. Control (dH₂O)-exposed brains served as the experimental control group for assessing baseline FosTRAP2-associated labeling.

Whole-brain image volumes were registered to the Allen CCFv3 (Wang et al., 2020) using SimpleElastix (Marstal et al., 2016). Registration incorporated anatomical information from the red, green, and blue fluorescence channels, using sequential translation, affine, and B-spline transformations adapted from our previous STPT workflow (Ramirez et al., 2025). These transformations were applied to the corresponding P0 and P3 probability maps, which were resampled to the 10-µm isotropic CCFv3 grid. Registered probability maps were quantified using custom Python scripts. For each brain, probability-map intensities were summed within CCFv3 anatomical masks. Left and right hemispheres were combined before volume normalization, yielding the bilateral signal density. A total of 316 bilateral CCFv3 summary regions were analyzed across Control, Snake1, and Tortoise1 groups. P0 and P3 signals, and their combined signal (P0+P3), were quantified separately. These measurements represent fluorescence-signal density. Regional densities were visualized using log₁₀(1 + density). Within-mouse AOB/MOB and AOB/MeA ratios were calculated from P0 densities using (AOB density + 1)/(MOB or MeA density + 1).

Each mouse was treated as an independent experimental unit. Regional signal densities were compared between Snake or Tortoise and Control using two-sided exact permutation tests, with FDR correction across 22 comparisons involving 11 selected regions. Effect sizes were quantified using Cohen’s *d*, with 95% confidence intervals estimated from 20,000 mouse-level bootstrap resamples. Exploratory relationships between regional FosTRAP2 signals and behavioral features were evaluated using group-adjusted Spearman correlations with exact within-group permutation tests.

### Statistical analysis

Statistical comparisons of state occupancy were performed separately for the two-chamber and multiple-choice assays. For the two-chamber assay (Cohort 1), PIMAP state occupancies were analyzed using one-way repeated-measures ANOVA with Greenhouse-Geisser correction for sphericity. Benjamini-Hochberg false discovery rate (FDR) correction was additionally applied across the 24 state-level omnibus tests. Pairwise comparisons were performed using two-sided paired *t* tests with Holm correction across all 15 pairwise odor comparisons within each state. For the multiple-choice assay, occupancies of five behavioral classes were analyzed using Friedman tests and two-sided paired Wilcoxon signed-rank tests, with Benjamini-Hochberg FDR correction across all six pairwise odor comparisons within each behavior. Statistical significance was defined as *p* < 0.05 for Friedman tests, Holm-adjusted *p* < 0.05 for paired *t* tests, and *q* < 0.05 for FDR-corrected comparisons.

## Results

### A 3D pose-based integrative machine-learning pipeline (PIMAP) identifies interpretable behavioral states during two-chamber risk assessment task

To quantify odor-evoked behavior across spatial, postural, and temporal dimensions, we upgraded our previous 2D behavioral-analysis framework (Wang et al., 2023) to a 3D Pose-based Integrative Machine-learning Analysis Pipeline (PIMAP; Fig. 1). Mice freely explored a two-chamber arena containing an exploration chamber with the odor stimulus and an adjacent hiding chamber. Behavior was recorded simultaneously from four cameras (Fig. 1A). FreiPose was used to reconstruct the 3D positions of 12 body landmarks distributed across the head, forelimbs, hindlimbs, and tail base (Schneider et al., 2022)(Fig. 1B), providing high-resolution measurements of body posture, movement, and spatial position relative to the odor source that were not readily accessible in our previous 2D workflow.

**Figure 1.**
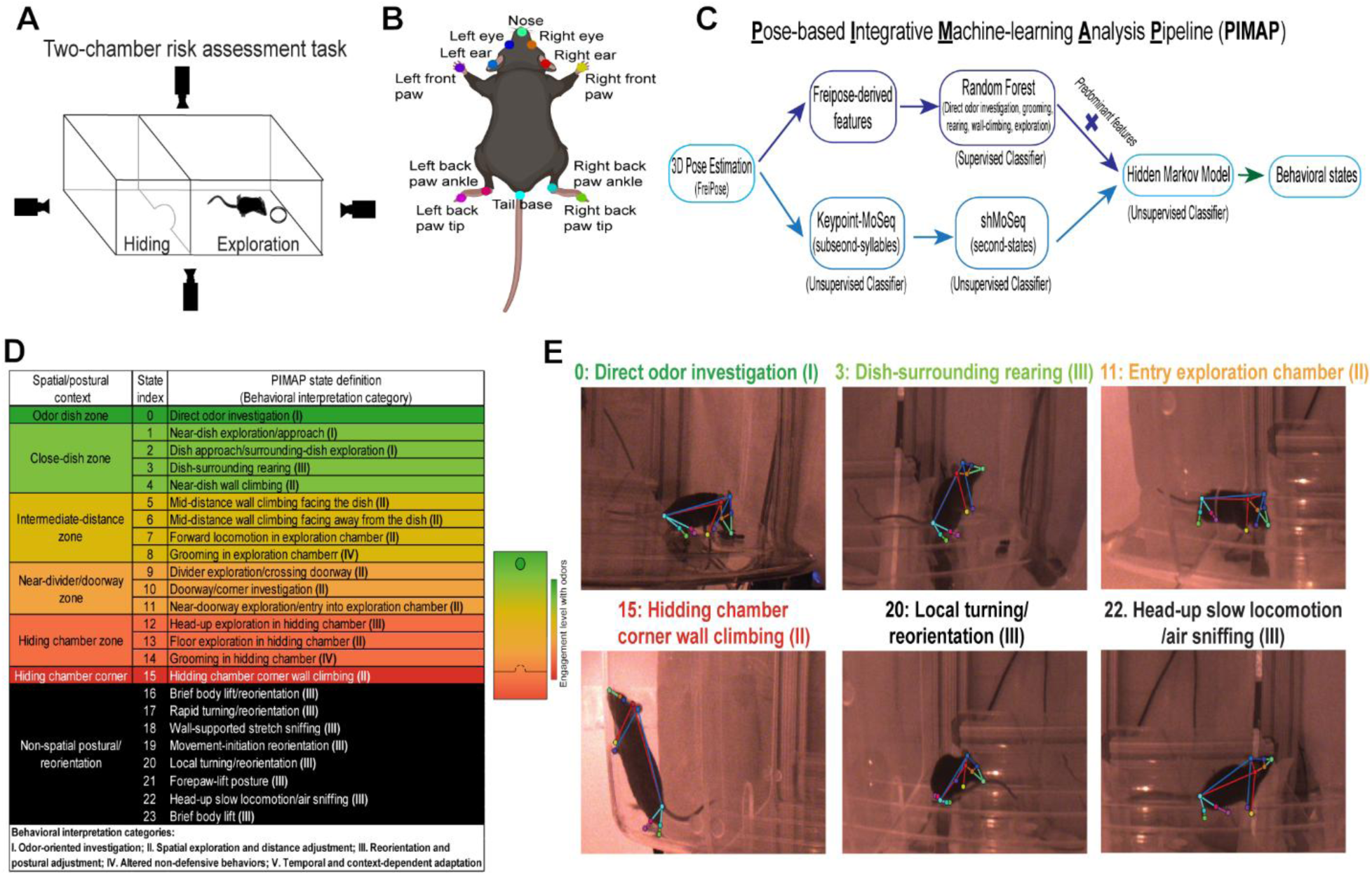
Overview of the two-chamber behavioral assay and pose-based integrative machine-learning analysis pipeline (PIMAP). (A) Schematic of the two-chamber risk-assessment task. Mice freely explored an arena containing a hiding chamber and an exploration chamber with the odor stimulus while behavior was recorded simultaneously from four camera views. (B) Twelve mouse body landmarks were used for 3D pose estimation through FreiPose, including the nose, left and right eyes, left and right ears, left and right front paws, left and right hind-paw ankles, left and right hind-paw tips, and tail base. (C) Architecture of the Pose-based Integrative Machine-learning Analysis Pipeline (PIMAP). Supervised Random Forest classifications, unsupervised Keypoint-MoSeq/shMoSeq representations, and odor-relative spatial information were integrated using a Gaussian hidden Markov model to generate the final PIMAP behavioral states. (D) Classification and interpretation of the 24 final PIMAP states (states 0-23). States were organized by spatial/postural context into odor-dish, close-dish, intermediate-distance, divider/doorway, hiding-chamber, hiding-chamber-corner, and non-spatial postural/reorientation groups. Color coding represents spatial engagement relative to the odor source, from direct odor investigation to more distant locations; non-spatial postural/reorientation states are indicated in black. Roman numerals indicate five behavioral interpretation categories: I, odor-oriented investigation; II, spatial exploration and distance adjustment; III, reorientation and postural adjustment; IV, altered non-defensive behaviors; and V, temporal and context-dependent adaptation. Individual signatures may contribute to more than one category. Category V represents the temporal dimension, including latency, active span, and changes in behavioral organization across the assay. (E) Representative video frames with reconstructed 3D poses illustrating selected PIMAP states: direct odor investigation (state 0), dish-surrounding rearing (state 3), entry into the exploration chamber (state 11), hiding-chamber-corner wall climbing (state 15), local turning/reorientation (state 20), and head-up slow locomotion/air sniffing (state 22).

The PIMAP integrated complementary supervised and unsupervised behavioral representations (Fig. 1C). Supervised Random Forest classification identified predefined, readily recognizable behaviors, while Keypoint-MoSeq and shMoSeq captured more subtle subsecond postural dynamics and longer-timescale behavioral organization, respectively (Weinreb et al., 2024; Weinreb et al., 2026). Outputs from the supervised and unsupervised models were integrated with odor-relative spatial information using a GHMM, providing a unified representation of behavioral states, including postural dynamics, movement patterns, and spatial context. The final two-chamber representation contained 24 PIMAP states, ranging from the odor-dish and close-dish zones to intermediate-distance, divider/doorway, hiding-chamber, and hiding-chamber-corner zones, together with non-spatial postural/reorientation states (Fig. 1D, E). To facilitate interpretation, individual states were color-coded according to their spatial engagement with the odor source, ranging from direct odor investigation to progressively more distant locations, with non-spatial postural/reorientation states represented separately. This organization provided an intuitive description of both the animal’s behavior and its spatial relationship to the odor source.

High-resolution behavioral analysis can reveal fine-scale postural and spatiotemporal patterns, and an explicit framework for their ethological interpretation is equally important (Datta et al., 2019; Pereira et al., 2020). Predator-associated cues can elicit highly variable behavioral responses that do not necessarily correspond to a single stereotyped defensive behavior (Apfelbach et al., 2005; Blanchard et al., 2011; Rosen et al., 2015). To provide a biological framework for subsequent behavior interpretation, PIMAP states and derived behavioral features were organized within five complementary categories informed by these previous studies: I, odor-oriented investigation; II, spatial exploration and distance adjustment; III, reorientation and postural adjustment; IV, altered non-defensive behaviors; and V, temporal and context-dependent organization. Categories I-IV primarily describe the behavioral states and their transitions, whereas Category V captures how these behaviors are organized and redistributed across time and context, including first-versus second-half dynamics, latency, active span, and temporal changes in state transitions. We therefore interpreted odor-associated responses at four complementary levels: (1) individual PIMAP states; (2) selected odor-associated behavioral signatures; (3) literature-informed behavioral categories; and (4) broader behavioral strategies inferred from combined spatial and temporal organization. This framework allowed us to characterize complex predator-associated behaviors beyond individual stereotyped responses while distinguishing directly measured behavioral features from their broader biological interpretation.

### Multidimensional behavioral organization reveals distinct odor-associated signatures during the two-chamber risk assessment task

In our Cohort 1 two-chamber assay, we first determined whether occupancy of individual PIMAP states was sufficient to distinguish behavioral responses across the six odor conditions (Fig. S1). The results showed that TMT produced the most prominent occupancy changes, with nearly absent direct odor investigation (state 0; I), reduced close-dish exploration (states 1, 2, and 4; I, II), and increased mid-distance wall climbing and divider/doorway exploration (states 6, 9, and 10; II; Fig. S1). These changes indicate a redistribution of behavior away from direct odor investigation toward more distal exploration, consistent with previously described aversive responses to TMT (Apfelbach et al., 2005; Rosen et al., 2015; Wang et al., 2023). In contrast, PIMAP state occupancy showed substantial overlap among natural odor conditions, indicating that occupancy alone captured limited differences among these responses.

Given this overlap, we next quantified behavioral organization using 1,634 features encompassing state occupancy, bout structure, movement, state transition probabilities, and temporal dynamics (Supplementary Table 1). LDA of this expanded feature space separated the six odor conditions (Fig. 2A). The discriminant axis with the largest target-versus-rest effect differed among odors: TMT on LD1 (|*d*|=11.90), Lizard on LD3 (|*d*|=4.35), Mouse and Tortoise on LD4 (|*d*|=2.52 and 2.99), and Snake on LD5 (|*d*|=4.10; Fig. 2B). The N_top behavioral features in these axes were used to identify candidate behavioral signatures, followed by Cohen’s *d* screening and multiclass logistic-regression classification (see Methods). Thus, natural odors that were poorly distinguished by individual-state occupancy showed greater separation in the multidimensional LDA space when spatial, sequential, movement, bout, and temporal information were considered together.

**Figure 2.**
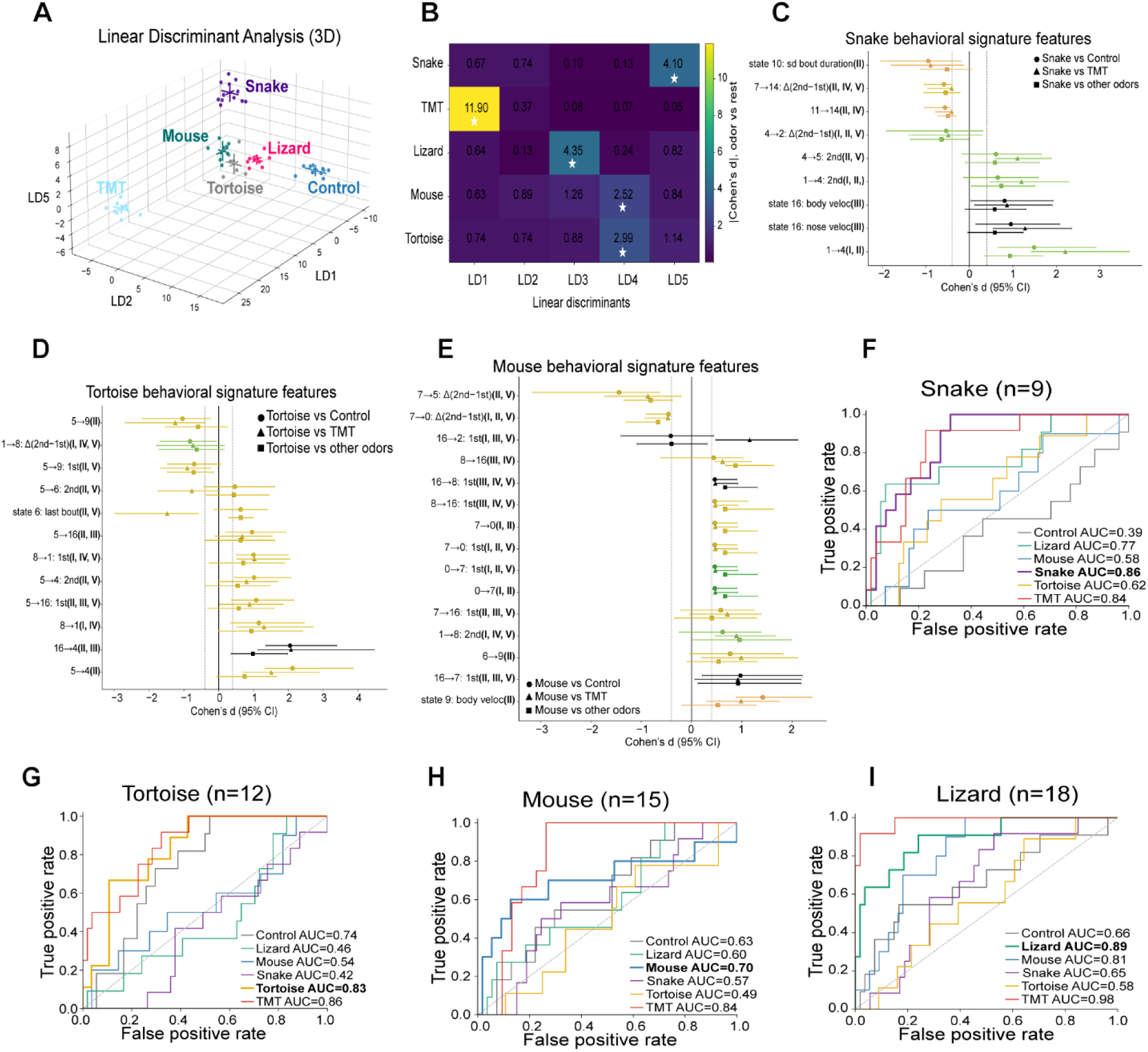
Multidimensional behavioral organization reveals distinct odor-associated signatures during two-chamber risk assessment. (A) Three-dimensional LDA projection of the 1,634-feature behavioral dataset for Control, Lizard, Mouse, Snake, Tortoise, and TMT conditions, shown along LD1, LD2, and LD5. Each point represents one analyzed mouse-odor session. Colored crosses indicate the group mean ± SD of individual animals’ LD scores. LDA was fitted using the complete Cohort 1 dataset, and the projections represent separation within the fitted data. (B) Absolute Cohen’s *d* comparing each target odor with the pooled remaining odors along LD1-LD5. Stars indicate the primary LD for each target, defined as the discriminant axis with the largest absolute target-versus-rest Cohen’s *d*: TMT, LD1; Lizard, LD3; Mouse and Tortoise, LD4; and Snake, LD5. (C-E) Cohen’s *d* and 95% confidence intervals for screened behavioral signatures identified for Snake (C, n=9 signatures), Tortoise (D, n=12 signatures), and Mouse (E, n=15 signatures). Candidate behavioral features were ranked by the largest absolute weights on the target-oriented primary LD and retained when the required comparator contrasts showed concordant directions and |*d*| ≥ 0.4 (see Methods). Circles indicate target versus Control, triangles indicate target versus TMT, and squares indicate target versus the pooled remaining natural odors. Signature colors indicate the spatial/postural context of the corresponding PIMAP state; for transition-probability signatures, color indicates the originating state. State transition features describe the probability of entering the destination state from the originating state, not the number of transition events. “1st” and “2nd” indicate signatures calculated during the first and second halves of the 5-min assay, respectively, and Δ(2nd−1st) identifies the difference between values measured during the second and first halves of the assay. Roman numerals indicate the behavioral interpretation categories defined in Fig. 1. (F-I) One-versus-rest ROC curves from multiclass logistic-regression models using the Snake (F), Tortoise (G), Mouse (H), and Lizard (I) behavioral signatures. Curves show classification performance for all six odor classes using the same target-derived signature set. ROC AUC was calculated from class probabilities averaged across four runs of mouse-grouped four-fold cross-validation. The n value in each ROC-panel title indicates the number of retained signatures. Cohort 1 included 12 WT mice (6 males and 6 females). Final analyzed sample sizes were Control, n=11; Lizard, n=11; Mouse, n=10; Snake, n=12; Tortoise, n=9; and TMT, n=12.

We next sought to identify the individual behavioral features that contributed most strongly to the LDA separation of behavioral responses across odor conditions. Post-LDA screening identified nine Snake-associated behavioral signatures, which yielded a target-class AUC of 0.86 (Fig. 2C, F). These signatures were dominated by spatial exploration and distance adjustment (II; 7/9), together with odor-oriented investigation (I), reorientation/postural adjustment (III), and altered non-defensive behavior (IV). Among the strongest signatures, Snake showed a greater likelihood of shifting from near-dish exploration to near-dish wall climbing (state 1→4) and increased body and nose velocities during brief body-lift/reorientation events (state 16). In contrast, bout-duration variability during doorway/corner investigation (state 10) was reduced, along with a lower probability of moving from near-doorway exploration to hiding-chamber grooming (state 11→14). Temporal signatures (V) included a higher probability of moving from near-dish wall climbing to mid-distance wall climbing during the second half (4→5: 2nd), while the tendency to move from forward locomotion to hiding-chamber grooming shifted less toward the second half than under other odors (7→14: Δ(2nd−1st)). Together, these signatures characterized Snake-associated behavioral responses involving near-dish investigation, repeated spatial adjustments, postural reorientation, and altered grooming-related transition probabilities, consistent with a multidimensional pattern of risk-assessment behavior rather than a simple shift toward approach or avoidance.

Tortoise, Mouse, and Lizard were also distinguished by multidimensional behavioral organization rather than changes in individual state occupancy. For Tortoise, 12 signatures yielded an AUC of 0.83, predominantly involving spatial exploration and distance adjustment (II; 9/12), including a greater likelihood of moving from mid-distance wall climbing while facing the dish to near-dish wall climbing (5→4) and a lower probability of shifting from the same mid-distance posture to divider exploration/doorway crossing during the first half (5→9: 1st) (Fig. 2D, G). Similarly, 15 Mouse signatures yielded an AUC of 0.70, primarily involving spatial exploration and distance adjustment (II; 10/15), together with postural reorientation (III). These included increased body velocity during divider exploration/doorway crossing (state 9: body veloc) and a smaller shift toward the latter half in the likelihood of moving from forward locomotion to mid-distance wall climbing while facing the dish, compared with other odor conditions (7→5: Δ(2nd−1st)) (Fig. 2E, H). For Lizard, 18 signatures yielded an AUC of 0.89, again dominated by spatial exploration and distance adjustment (II; 14/18), with additional investigation, postural, and grooming-related components (I, III, IV). Representative signatures included a greater likelihood of shifting from brief body lifting/reorientation to mid-distance wall climbing while facing the dish (16→5: 2nd) and a lower probability of moving from near-doorway exploration to divider exploration/doorway crossing (11→9: 2nd) during the second half (Fig. S2A; Fig. 2I). Together, these signatures revealed odor-associated differences in the spatial and temporal organization of behavioral sequences rather than changes in individual states alone.

Compared with natural fecal cues, TMT elicited a more pronounced redistribution of behavioral states, characterized by markedly reduced direct odor investigation. Post-LDA screening identified 50 TMT signatures, yielding a target-class AUC of 1.00 in the post-selection classification analysis (Fig. S2B, C). Among the 15 strongest displayed signatures, odor-oriented investigation (I; 12/15) was dominant, including large reductions in active span of direct odor investigation (state 0), last-bout timing, body and nose velocity, bout duration, and transition probabilities involving direct odor investigation. These reductions primarily reflected the near-complete absence of direct odor investigation rather than altered movement or bout duration during ongoing investigation. Additional signatures involved temporal organization (V; 7/15) and increased doorway/divider exploration (II; 2/15). Together, these TMT signatures reflect suppressed direct odor investigation and redistribution toward more distal behavioral states, consistent with its established aversive response (Rosen et al., 2015).

### Simultaneous presentation reveals distinct behavioral signatures across natural odors

To determine whether simultaneous exposure to multiple natural fecal odors elicited distinct behavioral patterns, we used a four-quadrant multiple-choice task containing Snake, Lizard, Mouse, and Tortoise odors (n=24 mice; Fig. 3A). We first compared occupancy of five matched PIMAP behavioral states within each odor quadrant (Fig. S3). Direct odor investigation was greater in the Snake quadrant than in the Lizard, Mouse, or Tortoise quadrant, and wall climbing was increased in the Snake quadrant relative to Lizard and Tortoise (Fig. S3C, E). Near-dish exploration was also higher for Snake than Lizard (Fig. S3A), whereas rearing did not differ significantly among odor quadrants (Fig. S3D). Grooming showed a different pattern, with lower grooming in the Snake quadrant than Lizard but greater grooming than Mouse and Tortoise (Fig. S3B). These results indicate increased Snake-associated odor investigation accompanied by odor-dependent changes in spatial and wall-oriented behavior, consistent with a multidimensional pattern of active risk assessment (Blanchard et al., 2011; Wang et al., 2023).

**Figure 3.**
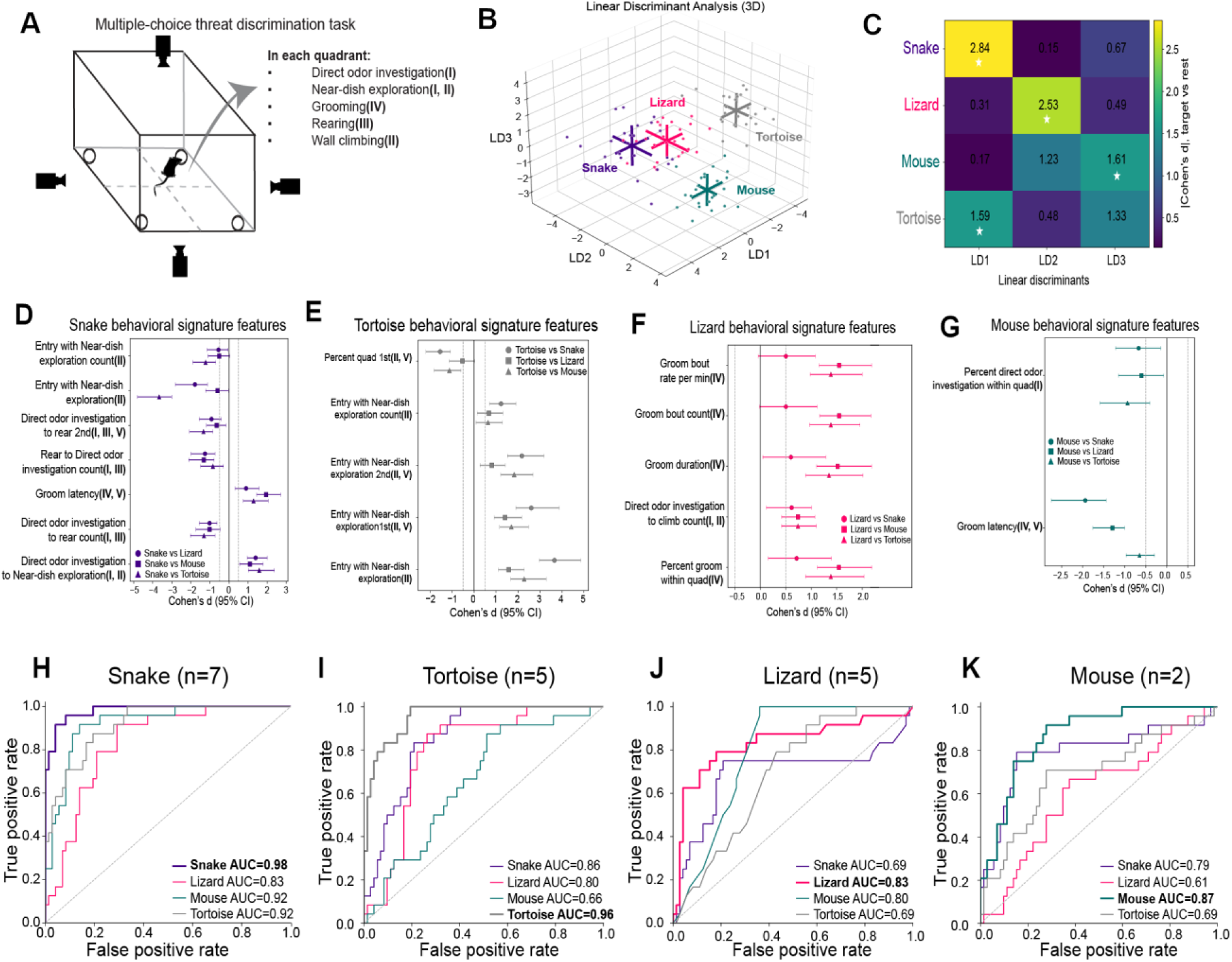
Multidimensional behavioral analysis identifies distinct signatures during simultaneous natural-odor presentation. (A) Schematic of the multiple-choice threat-discrimination task. Snake, Lizard, Mouse, and Tortoise odors were presented simultaneously in separate quadrants. PIMAP behavior states within each quadrant included direct odor investigation (I), near-dish exploration (I, II), grooming (IV), rearing (III), or wall climbing (II). Mice were habituated for two days with water-containing Petri dishes in all four corners before a single 20-min odor-exposure session. Odor locations were randomly assigned for the initial session and then held constant across mice to minimize cross-contamination during stimulus refreshing. (B) Three-dimensional LDA projection of the 144-feature behavioral dataset for the four odor conditions. Each point represents the behavioral response of one mouse within one odor quadrant. Colored crosses indicate the group mean ± SD of individual animals’ LD scores. (C) Absolute Cohen’s *d* comparing each target odor with the pooled remaining odors along LD1-LD3. Stars indicate the primary LD for each target, defined as the discriminant axis with the largest absolute target-versus-rest Cohen’s *d*: Snake, LD1; Lizard, LD2; Mouse, LD3; and Tortoise, LD1. (D-G) Cohen’s *d* and 95% confidence intervals for screened signatures identified for Snake (D, n=7 signatures), Tortoise (E, n=5 signatures), Lizard (F, n=5 signatures), and Mouse (G, n=2 signatures). Candidate behavioral features were ranked by absolute weight on each target’s primary LD. The 10 highest-ranked features were screened and retained as signatures when comparisons with each of the other three odors showed concordant directions and |d| ≥ 0.5. “Entry with [behavior]” refers to the behavior initiated upon entering an odor-associated quadrant; the corresponding count represents the number of such entries, and the percentage represents their proportion among all entries. These are distinct from transitions between behaviors. “1st” and “2nd” indicate signatures calculated during the first and second 10-min halves of the assay, respectively; Δ(2nd−1st) identifies the difference between the corresponding second- and first-half measurements. Roman numerals indicate the behavioral interpretation categories defined in Fig. 1. (H-K) One-versus-rest ROC curves from multiclass logistic-regression models using the Snake (H), Tortoise (I), Lizard (J), and Mouse (K) behavioral signatures. ROC AUC was calculated from class probabilities averaged across four runs of mouse-grouped four-fold cross-validation. Target-class AUCs were 0.98 for Snake, 0.96 for Tortoise, 0.83 for Lizard, and 0.87 for Mouse. The n value in each ROC-panel title indicates the number of retained signatures. The multiple-choice dataset included n=24 mice, with each animal contributing behavioral measurements for all four odor quadrants.

We next asked whether higher-dimensional behavioral organization further distinguished responses to four odors. Analysis of 144 behavioral features, encompassing quadrant occupancy, bout structure, quadrant-entry behavior, transition probabilities, and temporal dynamics, separated the four odor conditions by LDA (Fig. 3B; Supplementary Table 2). The strongest target-versus-rest separation occurred along LD1 for Snake (|*d*|=2.84), LD2 for Lizard (|*d*|=2.53), LD3 for Mouse (|*d*|=1.61), and LD1 for Tortoise (|*d*|=1.59; Fig. 3C). Post-LDA effect-size screening identified seven Snake signatures, yielding a target-class AUC of 0.98 (Fig. 3D, H). The Snake signatures most frequently involved direct odor investigation (I; 4/7), with additional contributions from spatial exploration (II), postural transitions (III), and grooming-related temporal organization (IV, V). The Snake quadrant showed reductions in both the number and percentage of entries beginning with exploration (Entry with near-dish exploration count; Entry with near-dish exploration). The probability of shifting from direct odor investigation to rearing during the second half was also reduced (Direct odor investigation to rearing 2nd), whereas the likelihood of moving from direct odor investigation to near-dish exploration (Direct odor investigation to near-dish exploration) and grooming latency (Grooming latency) were increased. Together with elevated Snake-directed investigation, these signatures suggest greater odor engagement accompanied by distinct patterns of quadrant entry, behavioral transitions, and delayed grooming, rather than generalized increases in quadrant exploration.

The other natural odors also showed distinct behavioral organization. For Tortoise, five signatures (AUC=0.96) were dominated by spatial exploration and distance adjustment (II; 5/5), including more quadrant entries beginning with near-dish exploration (Entry with near-dish exploration count), a greater percentage of quadrant entries beginning with near-dish exploration across the assay and in each half (Entry with near-dish exploration; Entry with near-dish exploration 1st; Entry with near-dish exploration 2nd), and reduced first-half quadrant occupancy (Percent quadrant 1st; V; Fig. 3E, I). The Lizard group showed five signatures (AUC=0.83), dominated by increased grooming bout rate, count, duration, and within-quadrant grooming (IV; 4/5), together with more transitions from direct odor investigation to climbing (I, II; Fig. 3F, J). The Mouse showed two signatures (AUC=0.87), comprising reduced within-quadrant direct odor investigation (I) and shorter grooming latency (IV, V; Fig. 3G, K).

### Reptile fecal cues recruit distinct vomeronasal sensory neuron populations

To determine whether the reptile fecal cues that produced distinct behavioral responses also generated distinguishable peripheral sensory representations, we performed volumetric Ca²⁺ imaging of VSNs using OCPI microscopy in OMP-Cre×Ai96 (OMP-GCaMP6s) vomeronasal epithelium (Holekamp et al., 2008; Turaga and Holy, 2012; Wong et al., 2020; Haran et al., 2026). Fecal extracts from two snake species, two lizard species, a tortoise, an iguana, and female mice elicited heterogeneous, stimulus-dependent responses across VSNs (Fig. 4A, B). Individual VSNs exhibited distinct response profiles, with representative neurons showing preferential responses to Snake1, Tortoise1, or female mouse feces, consistent with cue selectivity at the population level (Haran et al., 2026).

**Figure 4.**
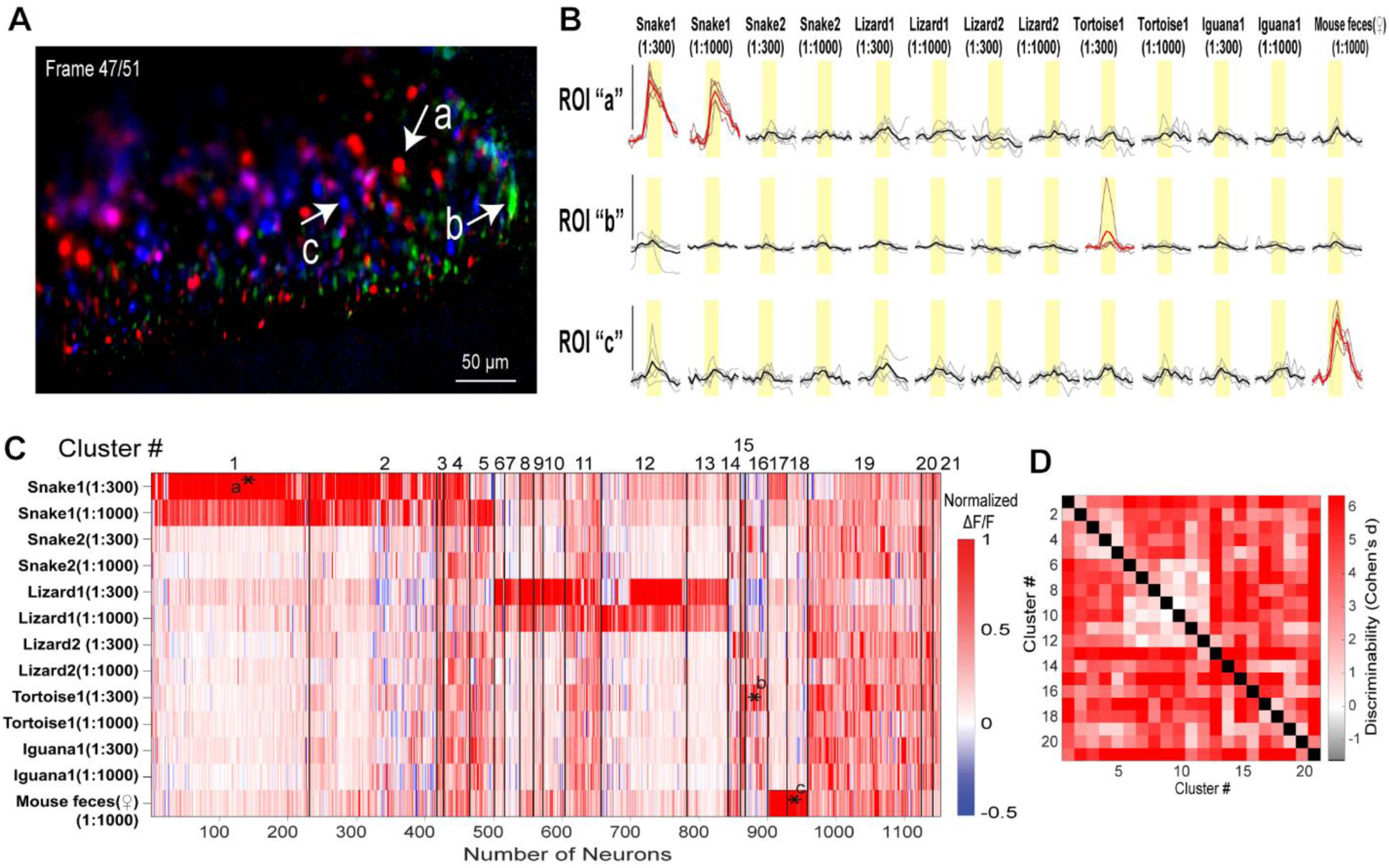
Reptile fecal extracts recruit distinct populations of mouse vomeronasal sensory neurons. (A) Representative pseudocolored image from volumetric OCPI Ca²⁺ imaging of vomeronasal epithelium expressing GCaMP6s in VSNs. Colored signals represent stimulus-evoked fluorescence responses, and arrows indicate representative VSN regions of interest (ROIs a-c) shown in B. The image represents one optical section from a 51 optical-section volumetric stack. Scale bar, 50 μm. (B) Across-trial ΔF/F responses of representative ROIs a-c to the fecal stimulus panel. Snake1 was collected from an eastern massasauga rattlesnake (*Sistrurus catenatus*), Snake2 from a Dumeril’s ground boa (*Acrantophis dumerili*), Lizard1 from a Henkel’s gecko (*Uroplatus henkeli*), Lizard2 from a great plated lizard (*Broadleysaurus major*), Tortoise1 from a pancake tortoise (*Malacochersus tornieri*), and Iguana1 from a green iguana (*Iguana iguana*); female mouse feces served as the conspecific comparison. Reptile fecal extracts were tested at 1:300 and 1:1,000 dilutions, and female mouse feces at 1:1,000. Stimuli were delivered for approximately 15 s in at least three randomized, interleaved repetitions. Thin traces indicate individual stimulus trials, bold traces indicate the across-trial mean, and yellow shading indicates stimulus presentation. Red traces identify stimulus-responsive VSNs according to the response criteria described in Methods (mean ΔF/F > 10% and *p* < 0.1 compared with Ringer’s control stimulation, two-tailed Student’s *t* test). (C) Clustered heatmap of normalized mean ΔF/F responses across the complete fecal stimulus panel. Each column represents an individual responsive VSN and each row represents one fecal stimulus and dilution. VSNs were organized into 21 functional response clusters using density-based merging (DBM) according to their response profiles; vertical black lines indicate cluster boundaries. The dataset included 1,152 analyzed VSN ROIs from 12 VNO tissues obtained from eight mice. Red indicates positive normalized ΔF/F, white indicates little or no response, and blue indicates negative ΔF/F. Asterisks labeled a-c indicate the clusters containing the representative ROIs shown in A and B. (D) Heatmap of pairwise discriminability (Cohen’s *d*) among the 21 functional VSN clusters, calculated using linear discriminant analysis. Higher Cohen’s *d* values (red) indicate greater separation between cluster response profiles, whereas lower values (lighter shading) indicate weaker separation. Black diagonal squares represent comparisons of each cluster with itself. Cohen’s *d* > 1.64 was used as the empirical criterion for satisfactory cluster separation, as described in Methods.

Population-level analysis further revealed heterogeneous and partially overlapping VSN responses to different fecal cues. We analyzed 1,152 VSN ROIs from 12 VNO tissues obtained from eight mice. Clustering of normalized response profiles identified 21 functional VSN clusters with distinct tuning across reptile samples and dilutions (Fig. 4C). Some clusters were strongly biased toward individual snake (e.g., clusters 1 and 2), lizard (e.g., clusters 6-8 and 13), tortoise/iguana (e.g., cluster 16), or female mouse cues (e.g., clusters 17 and 18), whereas others responded to multiple fecal extracts (e.g., cluster 19). Responses to the two samples within the snake and lizard groups were also not identical, indicating that VSN populations encoded information beyond broad reptile category alone. Pairwise linear discriminant analysis further revealed differences in the discriminability of these functional clusters, with many cluster pairs showing substantial separation in their response profiles (Fig. 4D). These distributed and partially selective response patterns are consistent with previous evidence that natural fecal chemosignals recruit heterogeneous VSN populations capable of representing species- and biological-state-related information (Wong et al., 2020; Haran et al., 2026). Together, these results demonstrate that predator-associated snake fecal cues generate distinct peripheral activity patterns within the VNO, providing sensory information that could contribute to subsequent odor discrimination and behavioral organization.

### Snake odor elicits a reproducible risk-assessment signature with widespread FosTRAP2 labeling

To determine how natural reptile odors are represented across central brain regions, we performed whole-brain STPT in FosTRAP2×Ai9 mice exposed to Control, Snake, or Tortoise odors. Following image quality control, 13 brains were retained for analysis (Control, n=5; Snake, n=5; Tortoise, n=3). We classified fluorescence signals into cell bodies (P0) and neuronal processes (P3) and quantified volume-normalized signal densities across 316 bilateral anatomical regions (Fig. 5A). Both signal classes showed heterogeneous distributions across brain regions and individual animals. Examination of 11 selected olfactory, limbic, hypothalamic, and hippocampal regions further revealed variation in regional labeling patterns (Fig. 5B). However, none of the regional comparisons remained significant after correction for multiple comparisons.

**Figure 5.**
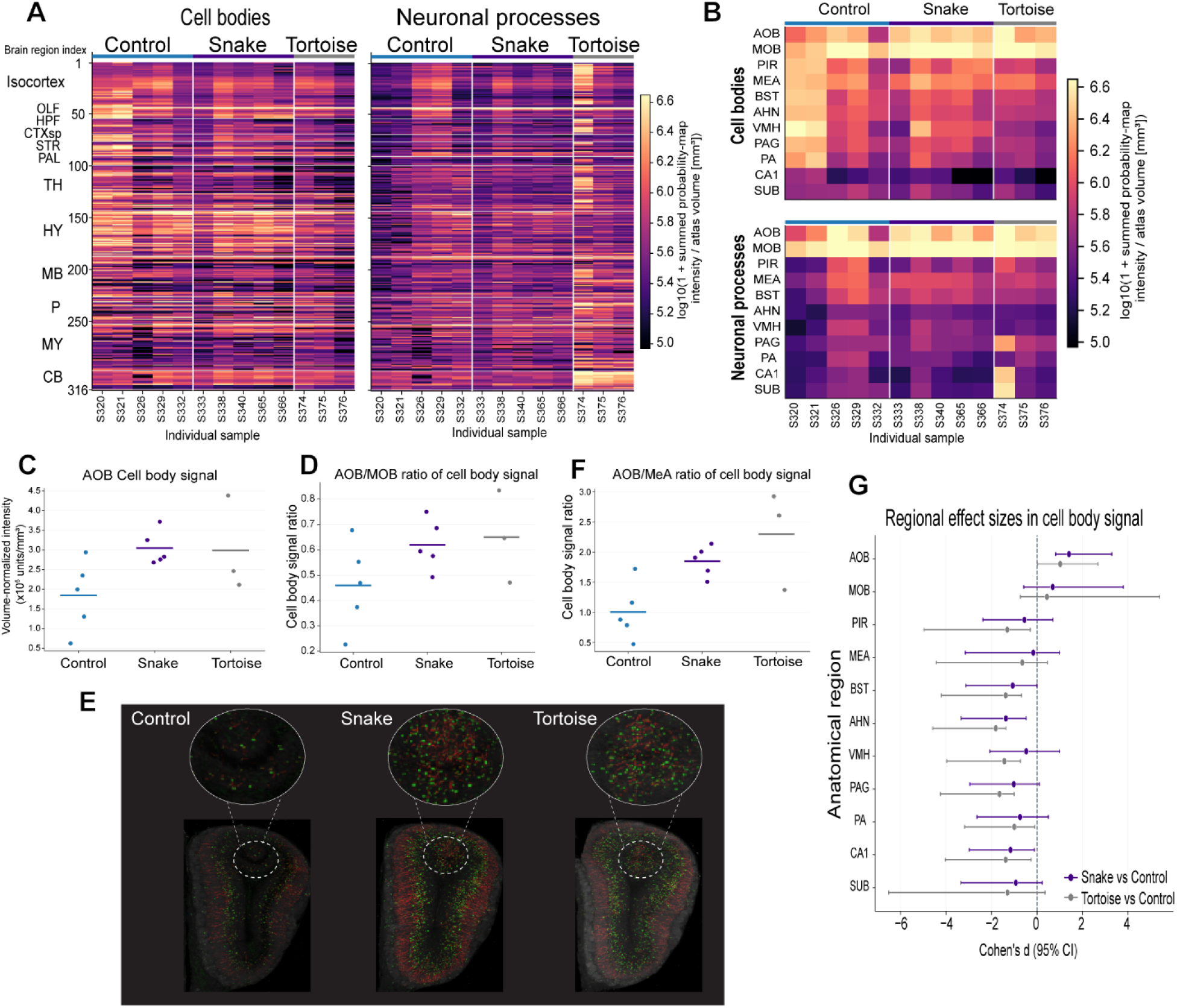
Whole-brain FosTRAP2 analysis characterizes regional cell-body- and neuronal-process-associated signals following natural odor exposure. (A) Heatmaps of cell-body (ilastik class 0; P0) and neuronal-process (class 3; P3) tdTomato classification-probability signals across 316 anatomically ordered Allen CCFv3 brain regions. Each row represents an anatomical region, and each column represents an individual mouse, grouped by Control (n=5), Snake (n=5), and Tortoise (n=3). For each region, signals from the left and right hemispheres were combined and normalized to the corresponding anatomical volume. Heatmaps display log10(1 + volume-normalized signal intensity), with identical color scales across the two signal classes. (B) Enlarged heatmaps of 11 anatomically selected regions associated with olfactory processing, limbic integration, and defensive behavior. Regional order, individual mice, normalization, and color scales are consistent with (A). (C) Volume-normalized AOB cell-body signal intensity across the three experimental conditions, expressed in millions of summed probability-map intensity units per mm³ (×10⁶ intensity units/mm³). (D) Within-mouse AOB/MOB ratio of cell-body signal densities (unitless). (E) Representative coronal fluorescence images of the AOB from Control, Snake, and Tortoise mice. Dashed circles indicate the regions enlarged in the corresponding upper insets. (F) Within-mouse AOB/MeA ratio of cell-body signal densities (unitless). Ratios in (D) and (F) were calculated from the original volume-normalized signals as (AOB+1)/(MOB+1) and (AOB+1)/(MeA+1), respectively. In (C), (D), and (F), each dot represents one mouse, and horizontal lines indicate group means. (G) Signed Cohen’s *d* effect sizes for cell-body signals in the 11 selected regions, comparing Snake versus Control (purple) and Tortoise versus Control (gray). Effect sizes were calculated from log10(1+volume-normalized signal intensity), and horizontal error bars indicate 95% confidence intervals. Positive and negative values indicate higher and lower signal intensities, respectively, relative to Control. AOB, accessory olfactory bulb; MOB, main olfactory bulb; PIR, piriform area; MeA, medial amygdala; BST, bed nuclei of the stria terminalis; AHN, anterior hypothalamic nucleus; VMH, ventromedial hypothalamic nucleus; PAG, periaqueductal gray; PA, posterior amygdalar nucleus; CA1, hippocampal field CA1; SUB, subiculum.

We next examined the AOB, the primary central target of VSNs. Mean AOB cell-body signal density was higher in Snake (3.05×10⁶ intensity units/mm³) and Tortoise (2.99×10⁶) than in Control (1.84×10⁶; Fig. 5C). Similarly, the mean AOB/MOB density ratios were 0.62, 0.65, and 0.46, respectively (Fig. 5D). Representative coronal images illustrated AOB labeling across the three conditions (Fig. 5E). These observations suggested relatively greater AOB labeling following both reptile odors, with comparable mean signals in the Snake and Tortoise groups. The corresponding AOB/MeA ratios were 1.85, 2.30, and 1.00 (Fig. 5F). Regional effect-size analysis showed positive AOB effects for both Snake versus Control (Cohen’s *d* =1.41) and Tortoise versus Control (d=1.02), whereas several other regions showed negative effect estimates (Fig. 5G). Nevertheless, no regional comparison survived FDR correction, and the observed pattern does not establish predator-specific regional recruitment.

To test whether this odor-exposure assay reproduced the behavioral responses observed in our previous experiments, we analyzed behavioral responses of these FosTRAP2×Ai9 mice. Similarly, LDA separated the three odor conditions in multidimensional behavioral space (Fig. S4A). Five Snake-associated signatures yielded a target-class AUC of 0.91 and predominantly involved spatial exploration and distance adjustment (II; 3/5) and odor-oriented investigation (I; 2/5; Fig. S4B, D). Snake exposure increased direct odor investigation (state 0) both overall and during the first half. During the second half, Snake exposure increased the likelihood of transitioning from hiding-chamber floor exploration to mid-distance wall climbing while facing away from the dish (state 13→6), while reducing transitions from the same exploratory state to hiding-chamber grooming (state 13→14). Transition probabilities from head-up slow locomotion/air sniffing to mid-distance wall climbing while facing away from the dish were also reduced (state 22→6). These signatures were consistent with increased odor source-directed investigation, along with altered spatial exploration and grooming-related transitions, as revealed in Cohort 1.

In contrast, nine Tortoise-associated signatures (AUC=0.85) primarily involved spatial exploration and distance adjustment (II; 5/9; Fig. S4C, E). Tortoise exposure reduced the likelihood of shifting from head-up hiding-chamber exploration to brief body lifting/reorientation (state 12→16) and late-session transitions from divider exploration to mid-distance wall climbing while facing the dish (state 9→5). Conversely, the probability of transitioning from head-up slow locomotion/air sniffing to mid-distance wall climbing while facing the dish increased (state 22→5). Tortoise also delayed dish approach and surrounding-dish exploration (state 2) and increased the median duration of hiding-chamber grooming bouts (state 14). Thus, Snake and Tortoise elicited distinct behavioral signatures despite their similar mean AOB cell-body signal densities.

Finally, we explored whether regional FosTRAP2 labeling was related to individual behavioral signatures in the retained mice for whole-brain imaging. We examined the relationship between direct odor investigation (state 0) during the first half of the 5-min behavioral recording (Fig. S4F) and labeling in the locus coeruleus (LC), a noradrenergic brainstem nucleus implicated in aversive and defensive responses (McCall et al., 2015; Gu et al., 2020). Group-adjusted Spearman analysis revealed a negative correlation between direct odor investigation and LC cell-body signal density (ρ=−0.88, exact within-group permutation p=0.0010; Fig. S4G). A similar relationship was observed after combining cell-body and neuronal-process signals (P0+P3; ρ=−0.91, p=0.0006; Fig. S4H). These findings suggest an association between LC labeling and odor-investigation behavior. This relationship indicates LC as a potential neural correlate of individual variation in odor-directed exploratory engagement.

### *Trpc2* loss differentially reorganizes odor-associated behavioral signatures

To determine whether odor-associated behavioral organization depends on TRPC2-mediated vomeronasal signaling, we compared WT and *Trpc2* KO mice in the two-chamber risk-assessment task. We first defined behavioral signatures using WT animals only. LDA separated the six odor conditions in multidimensional behavioral space (Fig. 6A), and WT-derived signatures classified Snake, Tortoise, and Mouse responses with target-class AUCs of 0.90, 0.96, and 1.00, respectively (Fig. 6B-D). TMT and Lizard signatures similarly showed strong within-WT classification, with AUCs of 1.00 and 0.89, respectively (Fig. S5A, B). Thus, WT mice exhibited distinct condition-associated behavioral signatures in this cohort.

**Figure 6.**
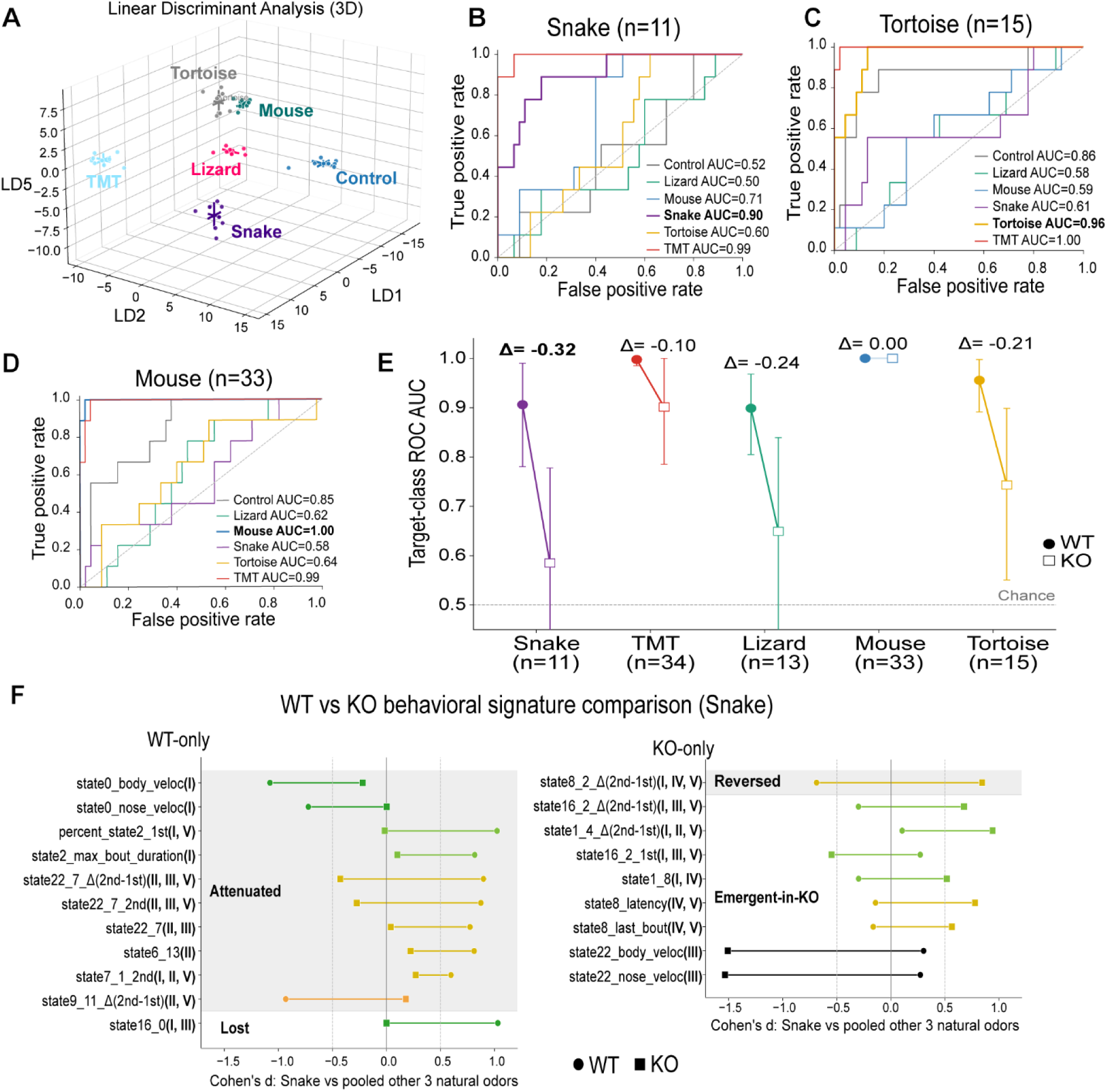
TRPC2 loss differentially reorganizes odor-associated behavioral signatures. (A) Three-dimensional LDA projection of WT behavioral features across Control, Lizard, Mouse, Snake, Tortoise, and TMT conditions. Each point represents one mouse-odor session. (B-D) One-versus-rest ROC curves from multiclass logistic regression using WT-derived Snake (B, n=11 signatures), Tortoise (C, n=15 signatures), and Mouse (D, n=33 signatures) signatures, with target-class AUCs of 0.90, 0.96, and 1.00, respectively. (E) Target-class AUCs of WT-derived signatures evaluated within WT mice and applied to KO observations using the original WT-selected signatures and WT-trained models without additional signature selection or model retraining. Filled circles indicate WT and open squares KO. Triangles indicate KO-minus-WT AUC. Changes were −0.32 for Snake, −0.10 for TMT, −0.24 for Lizard, 0.00 for Mouse, and −0.21 for Tortoise. The horizontal dashed line indicates chance-level AUC (0.5); error bars indicate 95% confidence intervals. (F) Comparison of the effect sizes between WT-derived (n=11) and KO-derived (n = 9) Snake behavioral signatures. All 11 WT and nine KO signatures were exclusive to their respective genotypes under the predefined selection criteria, with no shared signatures. Circles indicate WT Cohen’s *d*, squares KO Cohen’s *d*, and connecting lines differences between genotype-specific estimates. Effect sizes compare Snake with pooled Lizard, Mouse, and Tortoise within each genotype, excluding Control and TMT. State transition features describe the probability of entering the destination state from the originating state, not the number of transition events. “1st” and “2nd” indicate signatures calculated during the first and second halves of the 5-min assay, respectively, and Δ(2nd−1st) identifies the difference between values measured during the second and first halves of the assay. Shaded regions indicate signature changes in magnitude and direction, such as attenuated, lost, reversed, or emergent-in-KO effect-size patterns. Roman numerals indicate the behavioral interpretation categories defined in Fig. 1. (I-V). Signature colors indicate PIMAP spatial/postural context. Cohort 3 included nine WT and nine Trpc2 KO mice (six females, three males per genotype).

We next applied each WT-defined signature set and its corresponding WT-trained logistic regression model to *Trpc2* KO mice without additional signature selection or model retraining. The Snake signatures showed the largest reduction in classifier performance, with target-class AUC decreasing from 0.90 in WT to 0.58 in KO mice (Δ=−0.32; Fig. 6E). Lizard and Tortoise classification also decreased (Δ=−0.24 and −0.21), whereas TMT showed a smaller reduction (Δ=−0.10) and Mouse classification was preserved (Δ=0.00). These differences suggest that TRPC2 contributes to odor-associated behavioral organization, particularly during Snake exposure, with comparatively limited changes in TMT and Mouse classification performance following *Trpc2* deletion (Leypold et al., 2002; Stowers et al., 2002; Papes et al., 2010).

To examine genotype-associated differences in behavioral signatures, we compared signatures independently identified in WT and KO mice using the same effect-size contrast between each target odor and the pooled other natural odors (Fig. 6F). In WT mice, 11 Snake signatures predominantly involved spatial exploration and distance adjustment (II; 6/11) and odor-oriented investigation (I; 5/11), with additional reorientation/postural components (III). Snake-associated decreases in body and nose velocities during direct odor investigation (state 0) were less pronounced in KO mice, as was the increased occupancy of dish approach and surrounding-dish exploration during the first half of exposure (state 2: 1st). Temporal organization (V; 5/11) also differed between genotypes. During the second half of exposure, WT Snake responses showed a greater likelihood of shifting from head-up slow locomotion/air sniffing to forward locomotion than responses to the other natural odors, whereas the corresponding Snake-associated effect was attenuated or reversed in KO mice (state 22→7: 2nd; state 22→7: Δ(2nd−1st)). Similarly, the increased likelihood of moving from brief body lifting/reorientation to direct odor investigation in WT Snake responses was nearly absent in KO mice (state 16→0).

In contrast, nine KO Snake signatures primarily involved odor-oriented investigation (I; 5/9), with additional reorientation (III) and altered non-defensive behavior (IV). None of these signatures overlapped with the WT Snake signatures under the predefined selection criteria. Temporal signatures (V; 6/9) revealed distinct patterns of behavioral transitions involving dish-associated exploration, postural adjustments, and grooming. Notably, the tendency to shift from exploration-chamber grooming to dish-associated exploration was relatively more prominent later in the session in KO mice but showed the opposite odor-associated pattern in WT mice, compared with their respective responses to the other natural odors (state 8→2: Δ(2nd−1st)). KO Snake responses also showed a greater tendency toward transitions from brief body lifting/reorientation to dish-associated exploration (state 16→2: Δ(2nd−1st)) and from near-dish exploration to near-dish wall climbing (state 1→4: Δ(2nd−1st)) as the session progressed, relative to the other natural odors. Additional KO signatures involved transitions from near-dish exploration to exploration-chamber grooming (state 1→8) and altered timing of exploration-chamber grooming (state 8). Body and nose velocities during head-up slow locomotion/air sniffing (state 22) were also reduced in KO Snake responses relative to the other natural odors. Together, these findings suggest that *Trpc2* deletion alters the organization of Snake-associated behavior, shifting the dominant signatures from spatial exploration and distance adjustment in WT mice toward odor-oriented investigation and altered temporal sequencing in KO mice.

We further examined signatures identified independently within KO mice (Fig. S5C-G). Despite the reduced classifier performance of the WT-derived Snake signatures, the KO-derived Snake signature remained strongly discriminative within the KO dataset (AUC=0.97). KO-derived Tortoise, Mouse, Lizard, and TMT signatures yielded AUCs of 0.86, 1.00, 0.60, and 0.99, respectively. The WT Tortoise signature primarily involved spatial exploration and distance adjustment (II; 8/15) and odor-oriented investigation (I; 7/15), with 14 of 15 signatures showing attenuated effect-size contrasts in KO mice. The five KO-only Tortoise signatures involved temporal organization (V; 5/5), particularly transitions between exploratory and postural states (Fig. S5H). In contrast, 15 Mouse signatures were shared between WT and KO mice (15/33 WT; 15/43 KO), with many preserved or enhanced effects involving hiding-chamber exploration, postural transitions, and temporal organization (Fig. S6). Together, these findings suggest that *Trpc2* contributes to odor-associated behavioral organization in a stimulus-dependent manner, with Tortoise signatures shifting toward temporal organization while many Mouse-associated features remain shared across genotypes.

## Discussion

The AOS contributes to predator chemosensation and defensive behavior (Masini et al., 2010; Papes et al., 2010; Pérez-Gómez et al., 2015). Our findings extend this framework by revealing distinct patterns of sensory and behavioral responses to complex natural fecal cues. Across different experimental contexts, Snake-associated chemosignals elicited sustained odor investigation accompanied by coordinated changes in spatial exploration, postural reorientation, and temporal organization. Although natural odors engaged overlapping behavioral components, their responses were distinguished by fine-scale differences in postural adjustments and spatiotemporal organization that extended beyond simple attraction or avoidance. These findings extend our previous characterization of snake-associated risk assessment (Wang et al., 2023) and are consistent with the dynamic nature of predator-evoked defensive responses (Apfelbach et al., 2005; Blanchard et al., 2011; Takahashi, 2014). At the sensory periphery, natural reptile and mouse fecal extracts recruited partially overlapping but distinguishable populations of VSNs, supporting distributed representations of complex chemosignals (Doyle et al., 2016; Wong et al., 2020; Haran et al., 2026). Whole-brain FosTRAP2 analysis further revealed heterogeneous regional labeling, although no regional comparisons remained statistically significant. Furthermore, *Trpc2* deletion reduced the transfer of WT-derived Snake behavioral signatures to KO mice, while independently derived KO signatures retained substantial discriminative information, suggesting altered behavioral organization rather than a uniform loss of responsiveness. Together, these findings support a framework in which distributed peripheral sensory representations are associated with distinct patterns of risk-assessment behavior, with *Trpc2* contributing to their normal organization.

Behavioral responses to natural chemosignals involve multiple levels of organization, including individual-state occupancy and the spatial, sequential, and temporal relationships among states. Established behavioral measurements, including freezing, avoidance, and investigation duration, provide important indices of defensive and exploratory behavior, while analyses of their spatial and temporal relationships offer complementary information about behavioral organization (Gomez-Marin et al., 2014; Berman, 2018; Mathis and Mathis, 2020). To examine these complementary dimensions, we expanded our previous layered hybrid ML workflow (Wang et al., 2023) into PIMAP by integrating 3D pose estimation, supervised behavioral classification, unsupervised pose dynamics, and odor-relative spatial information within a common behavioral representation (Wiltschko et al., 2015; Bohnslav et al., 2021; Hsu and Yttri, 2021; Segalin et al., 2021; Goodwin et al., 2024; Weinreb et al., 2024; Weinreb et al., 2026). This integration enabled us to “zoom in” on fine-scale behavioral organization alongside state occupancy, including bout structure, movement, state transitions, and temporal dynamics. Importantly, although individual-state occupancy showed substantial overlap among natural odor conditions, these multidimensional signatures revealed differences in their spatial and temporal behavioral organization. These findings extend our previous hybrid analytical framework (Wang et al., 2023) and are consistent with the view that behavior is hierarchically organized across multiple spatial and temporal scales (Gomez-Marin et al., 2014; Berman, 2018; Weinreb et al., 2026). Thus, PIMAP offers a complementary, high-resolution characterization of natural chemosignal responses by integrating individual-state occupancy with postural and spatiotemporal features.

Identifying subtle behavioral differences among natural odors is important, but interpreting their biological significance remains equally challenging (Gomez-Marin et al., 2014; Berman, 2018). To address this challenge, we established a hierarchical interpretation framework connecting individual PIMAP states and derived signatures to literature-informed behavioral categories and broader behavioral strategies. To distinguish the behavioral components expressed from their temporal organization, we defined five complementary categories. Categories I-IV describe odor investigation, spatial exploration and distance adjustment, postural reorientation, and altered non-defensive behaviors, whereas Category V captures their temporal organization (Apfelbach et al., 2005; Blanchard et al., 2011). This framework allows us to “zoom out” to connect individual behavioral signatures to their broader biological organization. Across experimental contexts, Snake-associated responses involved sustained odor investigation and spatial adjustment, whereas Tortoise, Lizard, and Mouse responses showed different contributions of spatial exploration, grooming, and postural transitions. Together, these findings suggest that natural odors engage overlapping behavioral components but organize them into distinguishable patterns. Thus, identifying fine-scale behavioral differences and integrating them into biologically meaningful strategies represent complementary steps toward understanding complex risk-assessment behavior.

Our physiological experiments further showed that natural fecal cues recruited heterogeneous, stimulus-biased VSN populations, with several response clusters showing limited overlap across stimuli. This organization is consistent with previous studies demonstrating distributed and partially overlapping VSN representations of natural and chemically defined chemosignals (Ben-Shaul et al., 2010; Isogai et al., 2011; Turaga and Holy, 2012; Tolokh et al., 2013; Fu et al., 2015). Although predator-derived proteins can activate selective VSN populations and elicit defensive behavior (Papes et al., 2010), fecal bile acids recruit multiple VSN populations with distinct sensitivities and tuning profiles (Doyle et al., 2016; Wong et al., 2020). Recent studies further indicate that fecal bile acid patterns contain information associated with species, diet, and gut microbiome status (Haran et al., 2026). Our observation of both shared and odor-specific VSN responses, including differences between fecal samples within the same broad reptile category, suggests that peripheral representations may encode multiple biological signatures beyond predator identity alone. Thus, distributed VSN population activity provides a potential sensory basis for distinguishing complex natural chemosignals, although the specific ligands and neuronal populations contributing to the observed behavioral responses remain to be identified.

*Trpc2* signaling contributed to the organization of Snake-associated behavior, but its disruption did not eliminate odor-associated behavioral differentiation. Previous studies have established that *Trpc2* is important for vomeronasal transduction and associated social and defensive behaviors (Leypold et al., 2002; Stowers et al., 2002; Papes et al., 2010). Consistent with this role, the WT-derived Snake signature showed the largest reduction in classifier transfer to *Trpc2* KO mice (AUC 0.90 to 0.58), whereas an independently derived KO Snake signature remained discriminative within the KO dataset (AUC 0.97). Signature-level comparisons further revealed differences in odor investigation, spatial transitions, postural adjustment, and temporal organization, suggesting that *Trpc2* deletion alters the organization of WT-associated behavioral patterns rather than uniformly suppressing Snake-associated responses. Several mechanisms may account for the retained behavioral differentiation, including parallel olfactory pathways (Masini et al., 2010; Ferrero et al., 2011; Dewan et al., 2013; Takahashi, 2014; Pérez-Gómez et al., 2015), residual signaling in *Trpc2*-deficient VSNs (Zhang et al., 2010), and developmental compensation following constitutive deletion. Because *Trpc2* is also expressed in subsets of sensory neurons in the main olfactory epithelium, the behavioral effects of *Trpc2* deletion cannot be attributed unambiguously to VNO dysfunction alone (Omura and Mombaerts, 2014). Together, these findings suggest that *Trpc2* contributes to the normal organization of Snake-associated risk assessment, but is not the sole factor capable of supporting distinguishable behavioral responses.

Peripheral chemosensory representations are processed through central neural circuits that contribute to the organization of defensive behavior (Pérez-Gómez et al., 2015). The medial amygdala (MeA) receives accessory-olfactory input and communicates with hypothalamic and brainstem networks implicated in defensive behavior. Within these networks, the ventromedial hypothalamus (VMH), dorsal premammillary nucleus (PMd), and periaqueductal gray (PAG) contribute to the organization of predator-associated responses . Previous studies have demonstrated that predator-associated information from parallel olfactory pathways converges in hypothalamic circuits, where organized neuronal ensembles represent chemosensory information and predator-threat imminence (de Andrade Carvalho et al., 2020; Nguyen et al., 2024). However, how complex natural predator chemosignals recruit distributed brain networks and how these responses relate to multidimensional risk-assessment behavior remain unclear. To investigate central responses to natural odors, we used FosTRAP2 labeling and whole-brain STPT to characterize regional fluorescence signals. Snake and Tortoise showed numerically higher AOB cell-body signal densities than Control, although these differences were not statistically significant. Moreover, both groups showed similar mean AOB labeling despite distinct behavioral signatures, suggesting that AOB signal magnitude alone does not explain their behavioral organization. Exploratory analysis further showed that greater LC-associated labeling covaried with reduced direct odor investigation. Given the established role of LC noradrenergic signaling in stress-related arousal and aversive behavior (McCall et al., 2015; McCall et al., 2017), this relationship identifies a candidate brain-behavior association for future causal testing. Together, these findings suggest that reptile odors are associated with higher mean AOB labeling and heterogeneous neural responses across the brain, with some regional variations potentially related to risk-assessment behavior.

Several limitations define the scope of this study. Natural fecal extracts contain complex chemosignal mixtures, and the specific ligands responsible for the observed VSN and behavioral responses remain unresolved. Because reptile samples also differed in species, diet, and potentially physiological state, these factors cannot be independently evaluated in the present experiments. In addition, the fixed odor sequence in Cohorts 1 and 3 confounds odor identity with testing day and exposure history, while signature selection and parameter optimization preceding cross-validation may lead to optimistic classification estimates (Ambroise and McLachlan, 2002). Constitutive *Trpc2* deletion may also produce developmental adaptations and affect *Trpc2*-expressing neurons outside the VNO, limiting attribution of behavioral changes exclusively to vomeronasal signaling (Leypold et al., 2002; Stowers et al., 2002; Omura and Mombaerts, 2014). Finally, the present study does not establish a direct causal relationship between specific VSN populations and risk-assessment signatures. Addressing these limitations will be important for determining how natural predator chemosignals are detected and transformed into coordinated behavioral responses.

Together, our findings support a framework in which natural predator-associated chemosignals recruit distributed VSN populations and are associated with distinct patterns of risk-assessment behavior. By extending our previous hybrid ML approach, PIMAP enables fine-scale characterization of behavioral organization, while the biologically grounded interpretation framework connects individual signatures to coordinated behavioral strategies. Whole-brain FosTRAP2 analysis provides an initial description of regional neuronal labeling and exploratory relationships with behavioral responses. Disruption of *Trpc2* altered WT-associated Snake behavioral signatures without eliminating condition-associated behavioral differentiation, suggesting that vomeronasal signaling contributes to normal behavioral organization but is not its sole determinant. Overall, these findings establish complementary sensory, anatomical, and behavioral descriptions of natural predator chemosensation and provide a foundation for investigating how distributed peripheral representations are transformed through central neural circuits into coordinated risk-assessment behavior.

## Acknowledgments

We thank members of the Chemosensation and Social Learning Laboratory for helpful feedback. We especially thank Michael Mastrangelo for essential assistance with animal husbandry. We thank the Seneca Park Zoo veterinarian Christopher McKinney and other staff for providing reptile fecal samples (Project No. 20225). We appreciate the Whole Brain Microscopy Facility at UT Southwestern Medical Center for experimental support with whole-brain imaging.

## Author contributions

J.W. and J.P.M. designed the project, analyzed the data, and wrote the manuscript. J.W. collected reptile samples, conducted behavioral experiments, and analyzed the behavioral data. V.H. performed microscopy experiments, and J.W. and J.P.M. analyzed the microscopy data. J.W. performed the whole-brain microscopy experiments, and J.W. and J.P.M. analyzed whole-brain microscopy data.

---

**Figure S1.**
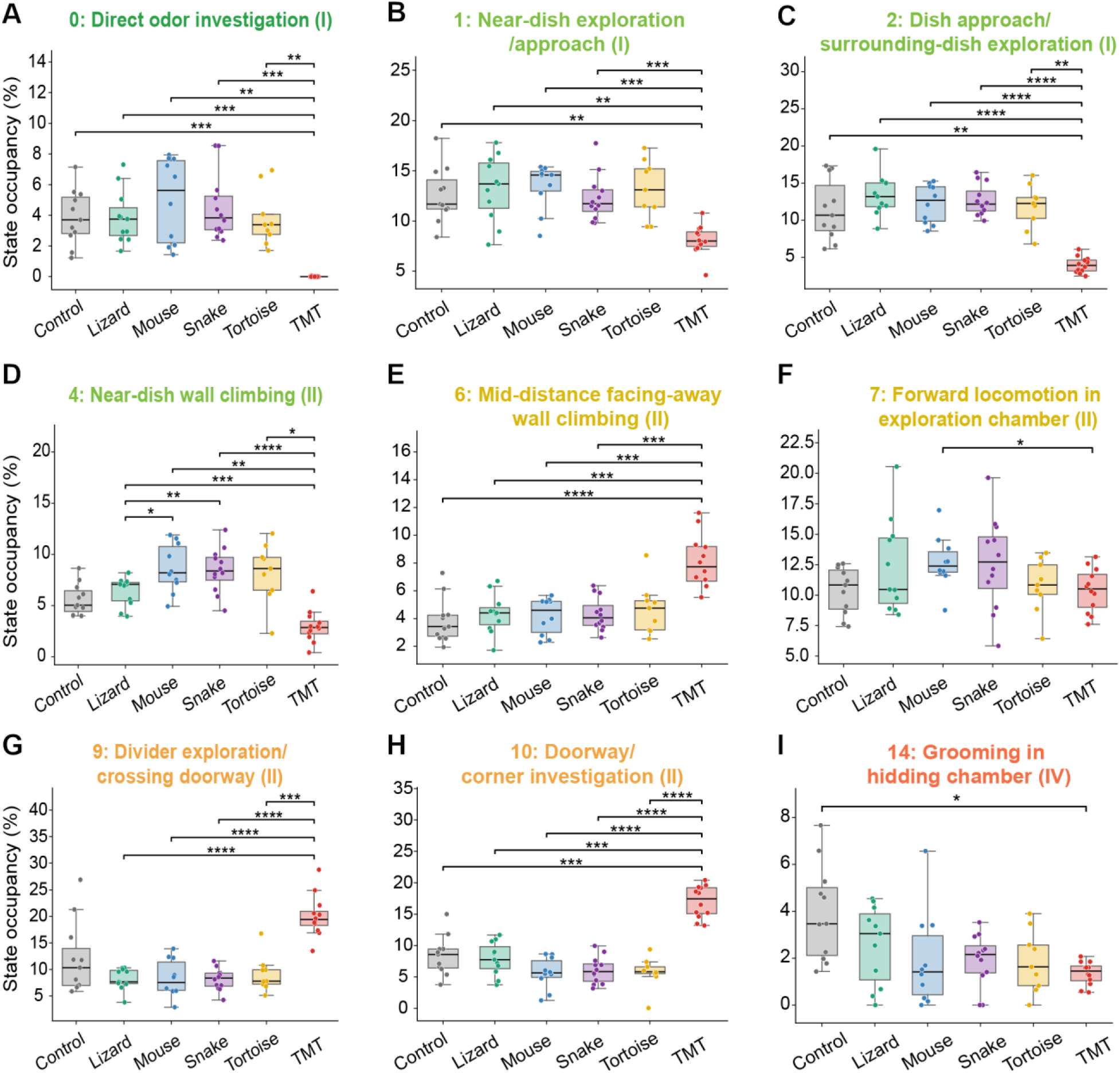
Individual-state occupancy primarily distinguishes TMT from other odor conditions in the two-chamber risk-assessment task. State occupancy was compared across Control, Lizard, Mouse, Snake, Tortoise, and TMT conditions for PIMAP states showing significant comparisons. (A) State 0, direct odor investigation (I). (B) State 1, near-dish exploration/approach (I). (C) State 2, dish approach/surrounding-dish exploration (I). (D) State 4, near-dish wall climbing (II). (E) State 6, mid-distance wall climbing facing away from the dish (II). (F) State 7, forward locomotion in the exploration chamber (II). (G) State 9, divider exploration/crossing doorway (II). (H) State 10, doorway/corner investigation (II). (I) State 14, grooming in the hiding chamber (IV). Roman numerals indicate the behavioral interpretation categories defined in Fig. 1. Occupancy represents the percentage of analyzed recording time assigned to each PIMAP state. Individual points represent animals, and box plots summarize the distribution within each odor condition. Overall odor effects were evaluated using one-way repeated-measures ANOVA with Greenhouse-Geisser correction when appropriate and Benjamini-Hochberg correction across the 24 state-level omnibus tests, followed by planned paired, two-sided t tests with Holm correction within each state. For the omnibus analysis, mice with missing odor sessions were excluded by complete-case analysis; individual paired comparisons used all mice with valid observations for the corresponding two conditions. Brackets indicate significant planned comparisons based on Holm-adjusted p values: *p < 0.05, **p < 0.01, ***p < 0.001, and ****p < 0.0001. Cohort 1 included 12 WT mice (6 males and 6 females). Final analyzed sample sizes were Control, n=11; Lizard, n=11; Mouse, n=10; Snake, n=12; Tortoise, n=9; and TMT, n=12. Matched-pair sample sizes ranged from 7 to 12 depending on the comparison.

**Figure S2.**
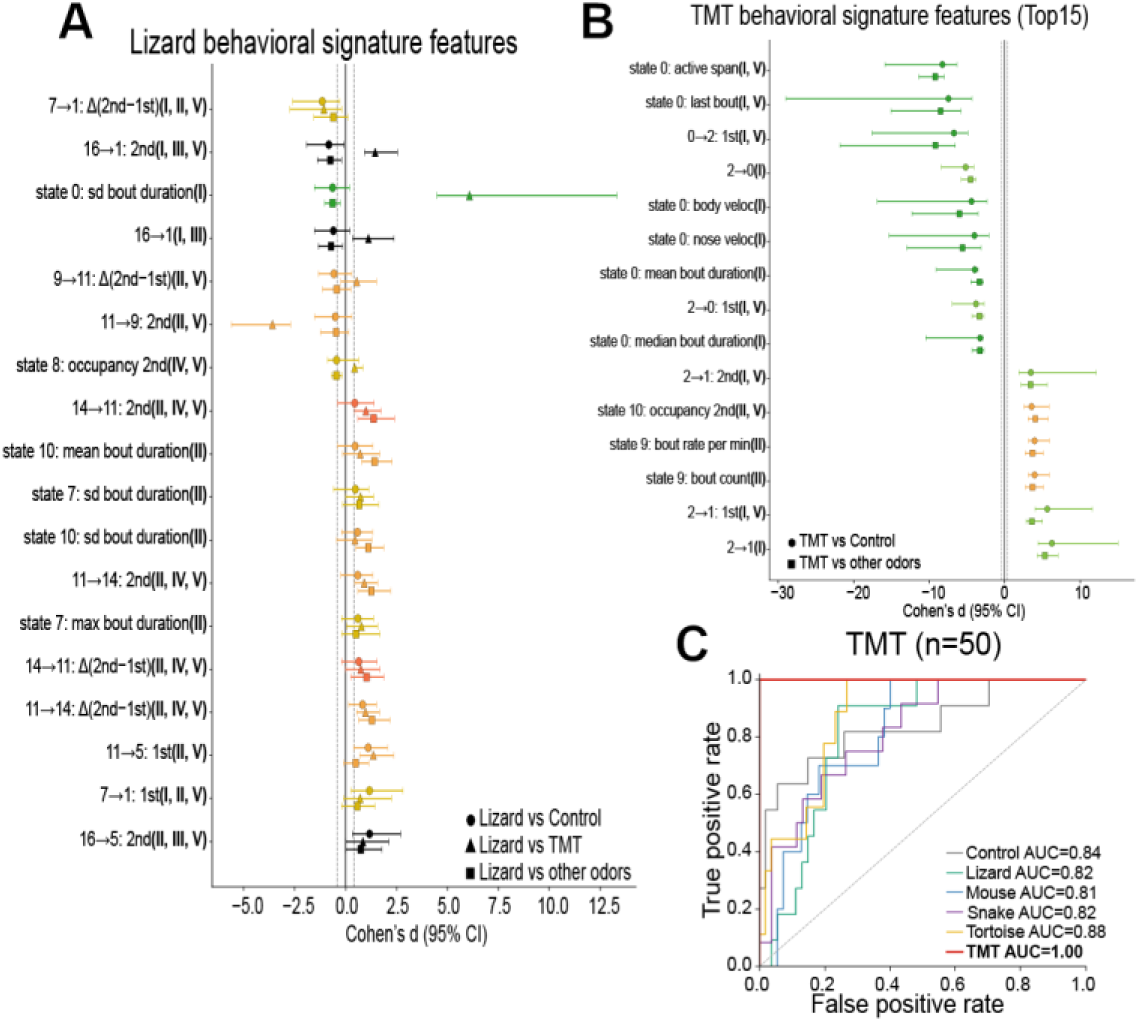
Lizard and TMT responses are characterized by distinct multidimensional behavioral signatures. (A) Cohen’s *d* and 95% confidence intervals for the 18 screened Lizard behavioral signatures. Circles indicate Lizard versus Control, triangles indicate Lizard versus TMT, and squares indicate Lizard versus the pooled remaining natural odors. (B) The 15 strongest of the 50 TMT behavioral signatures. Circles indicate TMT versus Control and squares indicate TMT versus the pooled remaining odor conditions. Signatures were identified from the largest absolute primary-LD weights followed by concordant effect-size screening (|*d*| ≥ 0.4). Signature colors indicate PIMAP spatial/postural context, and Roman numerals indicate the behavioral interpretation categories defined in Fig. 1. “1st” and “2nd” indicate signatures calculated during the first and second halves of the 5-min assay, respectively, and Δ(2nd−1st) identifies the difference between values measured during the second and first halves of the assay. (C) One-versus-rest ROC curves from post-selection multiclass logistic-regression classification using the complete 50 TMT signatures. ROC AUC was calculated from class probabilities averaged across four runs of mouse-grouped four-fold cross-validation. The n=50 in the panel title indicates the number of retained TMT signatures. Cohort 1 included 12 WT mice (6 males and 6 females). Final analyzed sample sizes were Control, n=11; Lizard, n=11; Mouse, n=10; Snake, n=12; Tortoise, n=9; and TMT, n=12.

**Figure S3.**
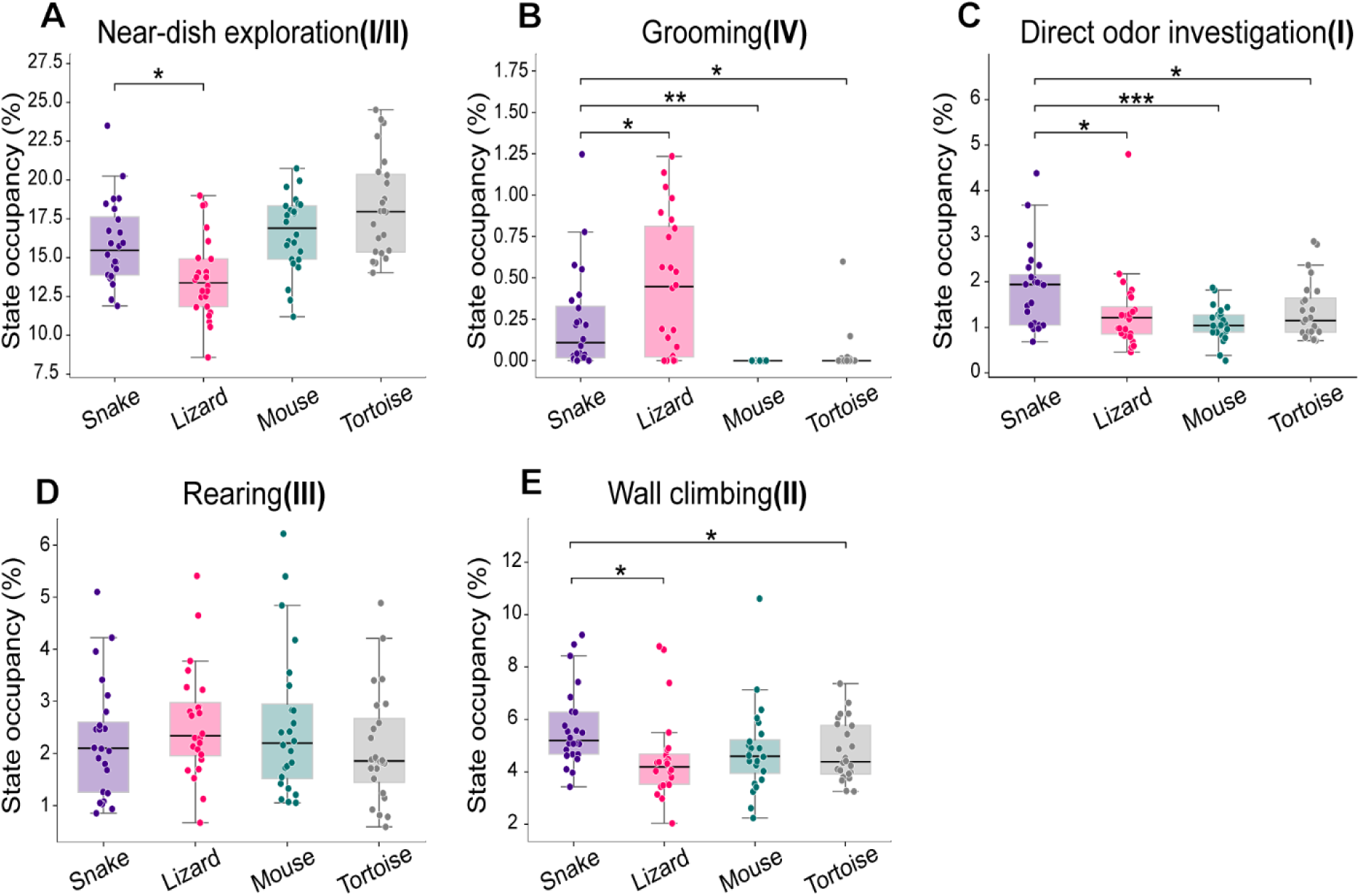
Behavioral responses differ among simultaneously presented natural odors. Occupancy of matched behaviors was compared among Snake, Lizard, Mouse, and Tortoise quadrants during the multiple-choice threat-discrimination task. (A) Near-dish exploration (I/II). (B) Grooming (IV). (C) Direct odor investigation (I). (D) Rearing (III). (E) Wall climbing (II). Roman numerals correspond to the behavioral interpretation categories defined in Fig. 1. Behavioral occupancy represents the percentage of the 20-min recording spent performing each specified behavior within the corresponding odor-associated quadrant. Individual points represent mice, and box plots summarize distributions across animals. Overall quadrant effects were evaluated using Friedman tests, followed by two-sided paired Wilcoxon signed-rank tests for all six odor-pair comparisons, with Benjamini-Hochberg FDR correction applied separately within each behavior. Only significant pairwise comparisons are shown; *q < 0.05, **q < 0.01, ***q < 0.001, based on FDR-adjusted q values. The same 24 mice contributed observations to all four odor-associated quadrants.

**Figure S4.**
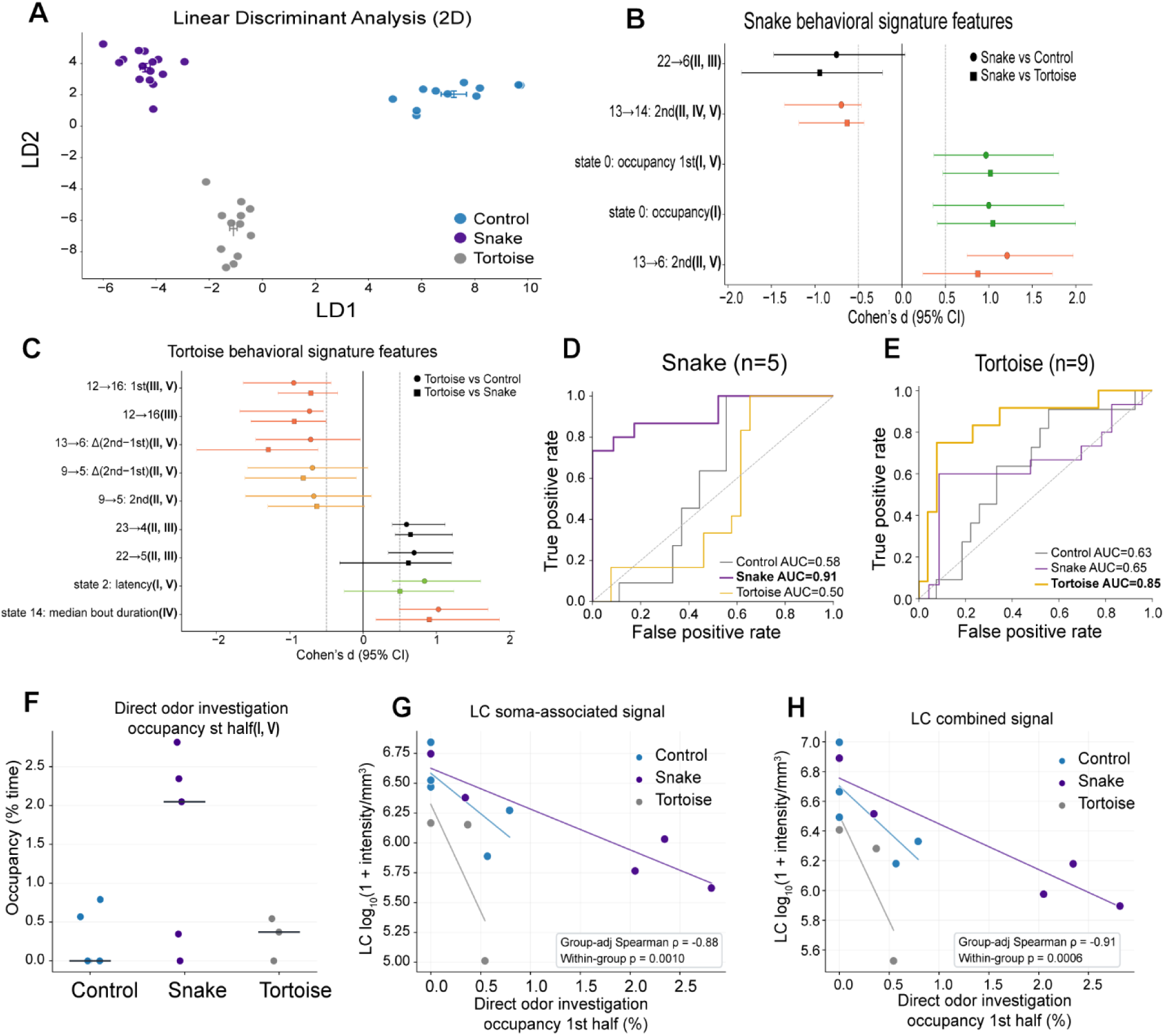
Odor-associated behavioral signatures and exploratory relationships between locus coeruleus signals and direct odor investigation in FosTRAP2 mice. (A) Two-dimensional LDA projection of multidimensional behavioral features for Control, Snake, and Tortoise conditions. Independent groups of FosTRAP2×Ai9 mice were habituated for two days and exposed to their assigned odor for 10 min on Day 3, following 4-hydroxytamoxifen (4-OHT) administration approximately 20 min before exposure. The first 5 min of exposure were used for behavioral analysis. Final analyzed sample sizes were Control, n=11 (6 males, 5 females); Snake, n=15 (7 males, 8 females); and Tortoise, n=12 (6 males, 6 females). Each point represents one mouse, and colored crosses indicate group means ± SD of individual LD scores. LDA projections represent separation within the fitted dataset. (B) Cohen’s *d* and 95% confidence intervals for the five screened Snake behavioral signatures. Circles indicate Snake versus Control, and squares indicate Snake versus Tortoise. (C) Cohen’s *d* and 95% confidence intervals for the nine screened Tortoise behavioral signatures. Circles indicate Tortoise versus Control, and squares indicate Tortoise versus Snake. Candidate features were ranked by their absolute weights on the target-specific primary LD and retained following concordant effect-size screening (|*d*| ≥ 0.5; see Methods). “1st” and “2nd” indicate features calculated during the first and second halves of the 5-min behavioral recording, respectively, and Δ(2nd−1st) indicates the difference between the two halves. Roman numerals indicate the behavioral interpretation categories defined in Fig. 1. (D, E) One-versus-rest ROC curves from post-selection multiclass logistic-regression models using Snake (D) and Tortoise (E) behavioral signatures. ROC AUC was calculated from class probabilities averaged across four runs of mouse-grouped four-fold cross-validation. The n values in the ROC-panel titles indicate the numbers of retained signatures. (F) Occupancy of direct odor investigation (PIMAP state 0) during the first half of the behavioral recording (0–2.5 min) in the 13 mice retained for whole-brain FosTRAP2 analysis (Control, n=5; Snake, n=5; Tortoise, n=3). Each dot represents one mouse, and horizontal lines indicate group medians. (G) Relationship between direct odor investigation occupancy during the first half and volume-normalized locus coeruleus (LC) cell-body signal (P0). (H) Relationship between the same behavioral measurement and combined LC cell-body and neuronal-process signals (P0 + P3). For (G) and (H), signals from both hemispheres were combined and normalized to the corresponding anatomical volume, followed by log10(1+signal intensity) transformation. Each point represents one mouse, and colored lines indicate descriptive within-group linear fits. Group-adjusted Spearman correlations were calculated after ranking both variables and centering their ranks within each experimental group. Two-sided *p* values were obtained using exact within-group permutations (86,400 permutations). LC, locus coeruleus.

**Figure S5.**
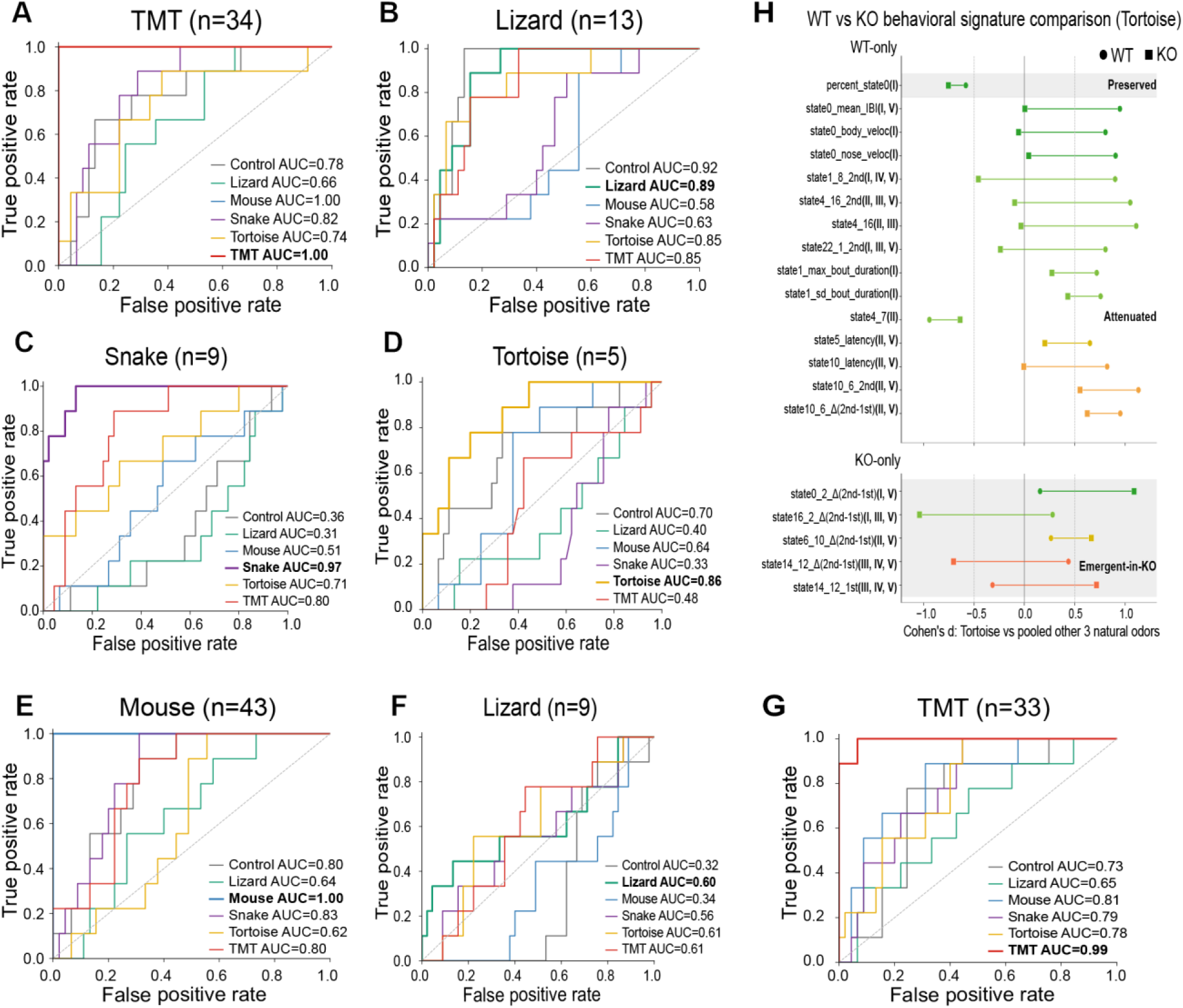
WT and Trpc2 KO mice exhibit distinct odor-associated behavioral signatures. (A,B) One-versus-rest ROC curves from post-selection multiclass logistic regression using WT-derived TMT (A, n=34 signatures) and Lizard (B, n=13 signatures) signatures, with target-class AUCs of 1.00 and 0.89, respectively. (C-G) ROC curves from independently identified KO signatures for Snake (C, n=9, AUC=0.97), Tortoise (D, n=5, AUC=0.86), Mouse (E, n=43, AUC=1.00), Lizard (F, n=9, AUC=0.60), and TMT (G, n=33, AUC=0.99). (H) Signature-level comparison of WT (n=15) and KO (n=5) Tortoise signatures. Independent signature screening identified 15 WT-only and five KO-only signatures, without shared signatures under the predefined selection criteria. Circles indicate WT Cohen’s *d*, squares KO Cohen’s *d*, and connecting lines indicate differences between estimates. Effect sizes compare Tortoise with pooled Snake, Lizard, and Mouse within each genotype, excluding Control and TMT. One WT-only signature showed a preserved effect-size pattern and 14 showed attenuation, whereas all five KO-only signatures were classified as emergent-in-KO. These classifications describe the relative magnitude and direction of Tortoise-associated effects in the two genotypes. Signature colors indicate PIMAP spatial/postural context. “1st” and “2nd” indicate signatures calculated during the first and second halves of the 5-min assay, respectively, and Δ(2nd−1st) identifies the difference between values measured during the second and first halves of the assay. Roman numerals indicate the behavioral interpretation categories defined in Fig. 1. Cohort 3 included nine WT and nine Trpc2 KO mice (six females, three males per genotype), with nine observations per odor condition and genotype.

**Figure S6.**
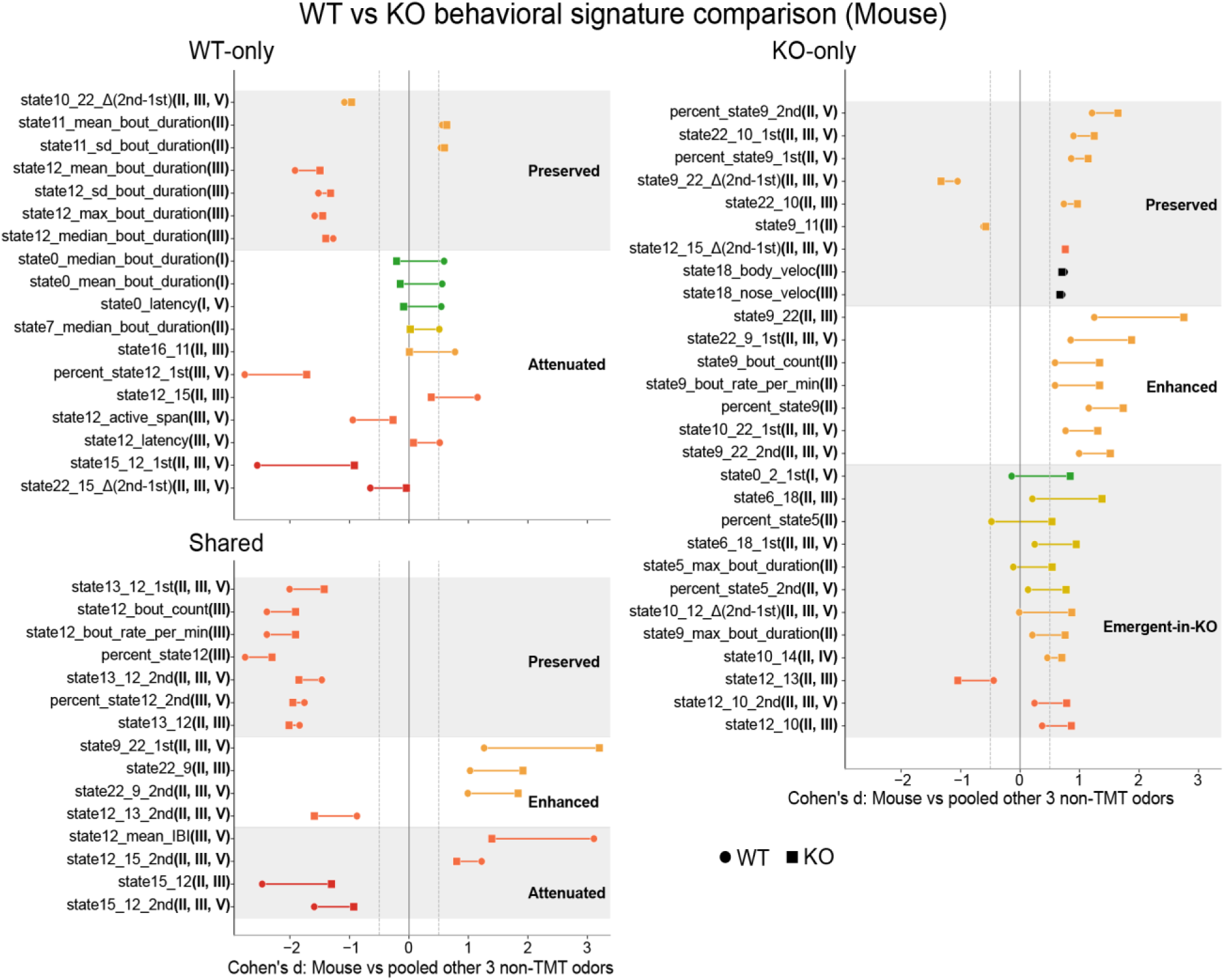
Mouse-associated behavioral organization is largely preserved but partially reorganized in *Trpc2* KO mice. Signature-level comparison of independently identified WT (n=33) and KO (n=43) Mouse signatures. Based on three-criterion signature selection, 18 signatures were classified as WT-only, 15 as Shared, and 28 as KO-only. Circles indicate WT Cohen’s *d*, squares KO Cohen’s *d*, and connecting lines differences between genotype-specific estimates. Effect sizes compare Mouse with pooled Snake, Lizard, and Tortoise within each genotype, excluding Control and TMT. Roman numerals indicate behavioral categories I-V, and signature colors indicate PIMAP spatial/postural context. Cohort 3 included nine WT and nine *Trpc2* KO mice (six females, three males per genotype), with nine observations per odor condition and genotype.

